# Through-scalp deep-brain stimulation in tether-free, naturally-behaving mice with widefield NIR-II illumination

**DOI:** 10.1101/2020.10.21.348037

**Authors:** Xiang Wu, Yuyan Jiang, Nicholas J. Rommelfanger, Rongkang Yin, Junlang Liu, Sa Cai, Wei Ren, Andrew Shin, Kyrstyn S. Ong, Kanyi Pu, Guosong Hong

## Abstract

Neural modulation techniques with electricity, light and other forms of energy have enabled the deconstruction of neural circuitry. One major challenge of existing neural modulation techniques is the invasive brain implants and the permanent skull attachment of an optical fiber for modulating neural activity in the deep brain. Here we report an implant-free and tether-free optical neuromodulation technique in deep-brain regions through the intact scalp with brain-penetrant second near-infrared (NIR-II) illumination. Macromolecular infrared nanotransducers for deep-brain stimulation (MINDS) demonstrate exceptional photothermal conversion efficiency of 71% at 1064 nm, the wavelength that minimizes light attenuation by the brain in the entire 400-1700 nm spectrum. Upon widefield 1064-nm illumination >50 cm above the mouse head at a low incident power density of 10 mW/mm^2^, deep-brain neurons are activated by MINDS-sensitized TRPV1 channels with minimal thermal damage. Our approach could open opportunities for simultaneous neuromodulation of multiple socially interacting animals by remotely irradiating NIR-II light to stimulate each subject individually.

## Introduction

Understanding complex neural circuitry and its correlation to specific behaviors requires spatially and temporally precise modulation of neuron subtypes in certain brain regions^1–3^. For decades, neural stimulation has been predominantly achieved with traditional electrical stimulation electrodes^4^. More recently, optogenetics has gained great popularity due to its rapid control of neural activities with visible light and its dissection of neural circuitry by selectively modulating specific neuron subtypes^1–3^. Alternatively, non-genetic optical neural interfaces have also been developed to allow neural stimulation with high spatiotemporal resolution^5–7^.

However, deep-brain neural modulation usually involves invasive implantation of stimulation electrodes and optical fibers due to the screening of electric fields and the scattering of light in the brain tissue^8–10^. Chronic brain implants lead to permanent damage to the brain tissue and overlying skull/scalp, while inducing a chronic immune response at the implant/tissue interface^10,11^. Furthermore, tethering the animal to an electrical wire or light source during behavioral studies leads to various deleterious consequences, especially for socially interacting animals (**Supplementary Table 1**)^3,12^. To mitigate these challenges, several novel methods have been demonstrated, including wireless optogenetic interfaces^3,13^, red-shifted opsins^14–18^, ultrasensitive opsins^18–20^, optogenetic antennas based on upconversion nanoparticles^21^, and sono-optogenetics based on mechanoluminescent nanoparticles^22^. Despite these recent advances, none of the existing optogenetic interfaces are able to altogether eliminate both the head tethering/fixing and the brain implants for deep-brain neural modulation in freely moving animals^13,21^. Magnetothermal neural modulation has been shown to free the animals from both head tethering and brain implants, yet it still requires a strong magnetic field inside a resonant coil that limits the free motion of the animals^23,24^. A noninvasive alternative is chemogenetics via peripherally or orally administered chemical actuators for selectively activating ligand receptors. However, chemogenetic neuromodulation has coarse temporal resolution due to the long residence time of chemical actuators *in vivo*^25,26^. Therefore, it is desirable to develop new methods that can modulate neural activity in the deep-brain regions of naturally behaving animals with high temporal resolution via a tether-free and implant-free interface.

Here we report macromolecular infrared nanotransducers for deep-brain stimulation (MINDS) in freely behaving animals through intact scalp and skull. MINDS can absorb light in one of the biological transparency windows, the second near-infrared window (NIR-II window, 1000-1700 nm; Fig. 1a)^27^, modulating neural activity via temperature-sensitive transient receptor potential (TRP) channels^28,29^. We demonstrate through-scalp deep-brain neuromodulation in tether-free and naturally-behaving animals with distant NIR-II illumination from >50 cm above the mouse head at a low incident power density of 10 mW/mm^2^. Our approach could open opportunities for simultaneous neuromodulation of multiple socially interacting animals by remotely irradiating NIR-II light to stimulate each subject individually.

**Fig. 1.**
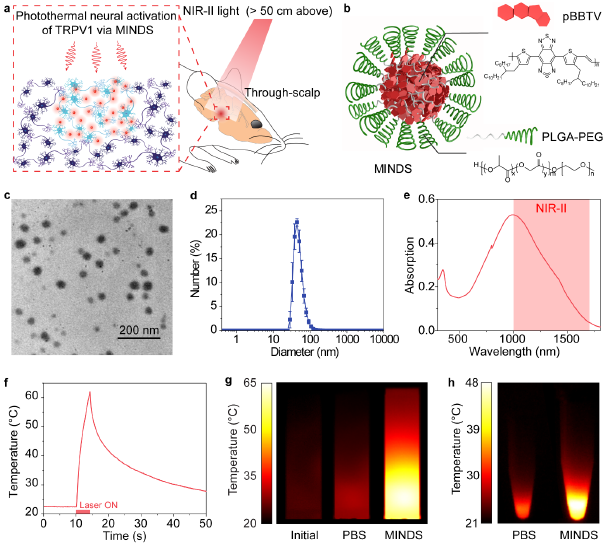
Efficient photothermal conversion of MINDS in the NIR-II window. (**a**) Schematic showing through-scalp neuromodulation with NIR-II illumination > 50 cm above the mouse head, which activates TRPV1 via the sensitization of MINDS (red circles). (**b**) Schematic showing the composition of MINDS, highlighting the pBBTV conjugated copolymers in the core (red hexagons) and the PLGA-PEG polymer comprising the shell (green spirals). (**c**) A representative TEM image of MINDS. (**d**) Distribution of the hydrodynamic diameters of MINDS revealed by DLS measurement. (**e**) Absorption spectrum of MINDS dispersion in PBS, showing strong absorption in the 1000-1700 nm NIR-II window (red shade). (**f**) Representative photothermal heating and cooling curve of 1.8 mg mL^−1^ MINDS dispersion, where the red bar indicates illumination with a 1064-nm laser at a power density of 10 mW mm^−2^. (**g**) Thermal images showing efficient photothermal heating of 25 μg mL^−1^ MINDS (right), in comparison with the same solution before NIR-II illumination (left) and PBS (middle). Both the PBS and MINDS images were taken when the temperature reached equilibrium after 30 min under 10 mW mm^−2^ 1064-nm illumination. (**h**) Thermal images of cell pellets incubated with PBS and MINDS, showing effective heating of cells with MINDS. Both images were taken when the temperature of the cell pellet reached equilibrium after 10 min under 10 mW mm^−2^ 1064-nm illumination.

## Results

### Choice of the NIR-II wavelength for deep-brain neuromodulation

We rationally chose 1064-nm light, since it offers the greatest tissue penetration by reduced scattering and similar, if not less, absorption in the brain tissue compared to visible light. In addition, 1064-nm light exhibits significantly lower water absorption than the longer-wavelength NIR-II spectrum beyond 1100 nm. We estimated the combined effect of scattering and absorption by calculating the brain extinction spectrum (**Supplementary Fig. 1**), which reveals that the 1064-nm wavelength is located near the minimum of the brain tissue attenuation spectrum in the entire 400-1700 nm range, which was validated by a previous report on light-tissue interaction^30^. Another advantage of 1064 nm, compared to other wavelengths around the attenuation spectral minimum, comes from its wide availability afforded by Nd:YAG lasers.

Owing to the minimum attenuation of 1064-nm light in the brain, it can penetrate to deeper brain regions than shorter wavelengths in the visible and traditional near-infrared (NIR-I, 700-900 nm) windows. Monte Carlo simulation reveals that 1064-nm NIR-II light can penetrate to a depth of at least 5 mm in the brain through the intact scalp and skull (**Supplementary Fig. 2**). Using the extinction coefficients of the scalp, skull, and the brain tissue, we estimated that 2.9% of the 1064-nm NIR-II light impinging on the scalp can reach a depth of 5 mm in the brain, in contrast to 1.4% of the 980-nm NIR-I light and 0.064% of the 635-nm red light for reaching the same depth (**Methods**). Therefore, although transcranial deep-brain optogenetics has been reported, existing techniques operated in the visible and NIR-I windows still require skull-tethered optical fibers to deliver light at a high incident power density to compensate for the loss in the brain^18,21^.

### Design principles and photothermal performance of MINDS

MINDS were designed with a π-conjugated semiconducting polymer core of pBBTV (poly(benzobisthiadiazole-*alt*-vinylene); **Supplementary Fig. 3&4**) to afford efficient absorption of 1064-nm light. This polymer core was coated with an amphiphilic and FDA-approved polymer shell, poly(lactide-co-glycolide)-*b*-poly(ethylene glycol) (PLGA-PEG; Fig. 1b) to afford water solubility and biocompatibility. MINDS have an average diameter of ~40 nm, as revealed by transmission electron microscopy (TEM, Fig. 1c) and dynamic light scattering (DLS, Fig. 1d).

MINDS have the following four key design principles that make them efficient NIR-II sensitizers to activate TRPV1 channels *in vivo*. First, MINDS are strong NIR-II light absorbers: the pBBTV core of the MINDS has strong absorption in the > 1000 nm NIR-II window (Fig. 1e) via adjustment of the highest occupied and the lowest unoccupied molecular orbitals (HOMO-LUMO) of the donor and acceptor units. Second, MINDS are efficient heat generators upon NIR-II illumination: under continuous 1064-nm illumination at 10 mW mm^−2^, 1.8 mg mL^−1^ MINDS solution reached 39 °C within merely 1.1 s (Fig. 1f&g). Specifically, the photothermal conversion efficiency of MINDS at 1064 nm was measured to be 71% (**Methods**), which represents one of the highest photothermal conversion efficiencies in the NIR-II window (**Supplementary Table 2**). Continuous 1064-nm illumination of human embryonic kidney (HEK) 293T cell pellets after incubation with MINDS resulted in significantly greater heating than the control group without MINDS (Fig. 1h). Third, MINDS remain stable in normal physiological condition and demonstrate superior photostability upon repeated NIR-II illumination. The structural stability of MINDS has been confirmed by the minimal change in either optical absorption or the hydrodynamic diameter over 7 days of incubation in phosphate buffered saline (PBS) and cell culture medium at 37 °C (**Supplementary Fig. 5**). Furthermore, the *in vivo* photostability of MINDS was proved by negligible variation in the photothermal performance for more than 70 heating/cooling cycles over 1 h (**Supplementary Fig. 6**). Fourth, the organic semiconducting polymer core and the PLGA-PEG shell render MINDS biocompatible. Compared to inorganic gold nanoparticles, organic semiconducting polymers have been reported to afford low toxicity and rapid biodegradation^31^. Primary neuron culture showed no reduction in cell viability after 24 h of incubation with MINDS up to a concentration of 3.6 mg mL^−1^ (**Supplementary Fig. 7**), while only a minimal chronic immune response was observed in the brain tissue near the injection sites of MINDS when compared with the carrier injection (**Supplementary Fig. 8**).

The superior photothermal performance of MINDS allows them to transfer heat to ectopically expressed TRP channels in neurons. Infrared-sensitive species such as venomous pit vipers sense long-wavelength radiation (750 nm – 1 mm) using TRP channels in a specialized organ^32^. More recently, ectopic expression of TRPV1 channels in mouse retinal cones endows vision in the infrared via the resonant absorption of gold nanorods^28^. Based on these results, we hypothesize that ectopic expression of TRPV1 channels in the mouse brain coupled with 1064-nm absorbing MINDS can achieve deep-brain neuromodulation with brain-penetrant NIR-II light. Several previous reports have used physical targeting of heat-producing nanoparticles to TRP channels and cell membranes^23,28,33^. However, we used untargeted MINDS for activating TRPV1 under NIR-II illumination in the following experiments, because the temporal dynamics of TRPV1 activation precludes the need for physically binding MINDS to TRPV1 channels. The fastest reported time constant of TRPV1 activation is 5 ms, achieved under a high laser power density of 1.0×10^6^ mW mm^−2^ to drive rapid temperature increase (**Supplementary Table 3**)^34^. Based on the Fourier heat equation, a thermal diffusion length of ~50 μm is expected over this timescale, much greater than the size of a neuron and its membrane. Similarly, a recent study rejects the existence of nanoscale heat confinement on the surface of particles^35^. Therefore, we reason that untargeted MINDS can provide sufficient heat diffusion to activate TRPV1 channels for both *in vitro* and *in vivo* experiments below.

### *In vitro* NIR-II photothermal activation of TRPV1 with MINDS

We then asked if focused NIR-II illumination was able to activate TRPV1 in the presence of MINDS. TRPV1 functions as a nonselective cation channel with a high permeability to calcium when activated^36^. Therefore, we used calcium imaging to demonstrate NIR-II activation of TRPV1 channels in HEK293T cells (Fig. 2a), which were transfected with pAAV-CMV-TRPV1-mCherry plasmids. Dynamic calcium imaging upon 1040-nm NIR-II illumination (parameters summarized in **Supplementary Table 3** and **Methods**) revealed several key findings.

**Fig. 2.**
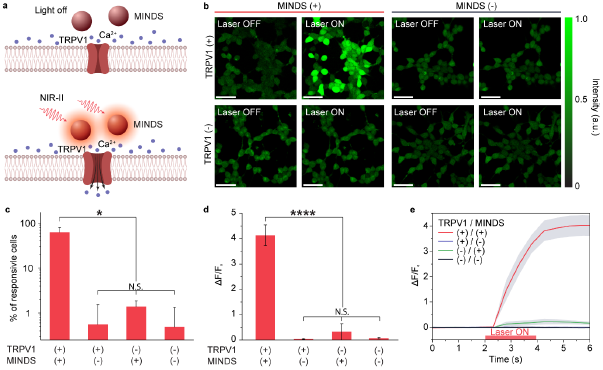
NIR-II photothermal activation of MINDS-sensitized TRPV1 *in vitro*. (**a**) Schematic showing NIR-II photothermal activation of TRPV1 expressed in the plasma membrane of cells mediated by MINDS. (**b**) Calcium imaging of HEK293T cells under different experimental conditions. A calcium signal increase is only seen when the NIR-II laser illuminates TRPV1+/MINDS+ cells. (**c**) Population study of 874 cells from twelve trials (three trials per group) in which the percentage of responsive cells is compared between different groups. A cell is defined responsive if its calcium signal rises over 50% of original intensity within 5 s of NIR-II illumination (i.e., Δ*F*/*F*_0_ > 50%)^24^. Error bars indicate standard deviation (SD). (**d**) Statistical analysis of calcium signal changes for different groups of cells (*n* = 15 cells for each group), shown as the ratio of maximum calcium signal change after NIR-II illumination over the original calcium signal before NIR-II illumination. In **c** & **d**, a statistically significant difference is found for TRPV1+/MINDS+ cells vs. all other conditions (*, *P* < 0.05; ****, *P* < 0.0001). There is no statistically significant difference in pairwise comparisons between TRPV1+/MINDS–, TRPV1–/MINDS+, and TRPV1–/MINDS– cells (N.S., not significant). Error bars indicate SD. (**e**) Temporal dynamics of the calcium signal for different groups of cells, showing an average latency time of 0.9 ± 0.2 s for calcium signal to increase over 3 SD of baseline and 1.1 ± 0.2 s to increase to 50% of the maximum. Shades indicate ±1 SD from *n* = 15 cells.

First, the specificity of calcium signal increase in MINDS+ and TRPV1+ cells suggests that MINDS-sensitized TRPV1 channels are activated under NIR-II illumination. 62.5 ± 17.3% of MINDS+/TRPV1+ cells responded to NIR-II stimulation with a more than 4-fold increase in calcium fluorescence intensity. In contrast, only < 1.5% of the cells in any control group showed a similar response within 5 s of NIR-II illumination, indicating the necessity of both MINDS and TRPV1 to increase the intracellular calcium concentration by NIR-II (Fig. 2b-d). We then proved that the observed calcium transients were neither from intracellular release of Ca^2+^ ions from endoplasmic reticulum calcium stores^37^ nor membrane capacitive current^5,38,39^. Specifically, TRPV1– cells showed no increase in calcium fluorescence even in the presence of MINDS upon NIR-II illumination (Fig. 2b-d). Furthermore, the cell specificity for NIR-II activation was confirmed by the colocalization of calcium signal increase and TRPV1 expression (**Supplementary Fig. 9**). In addition, MINDS– cells showed little calcium increase, suggesting that heat transfer to ectopically expressed TRPV1 channels via direct NIR-II illumination is inefficient and requires higher intensities of photon flux that would damage the neural tissue^28^. Therefore, the strong absorption of NIR-II light by MINDS sensitizes TRPV1 to low-intensity NIR-II light for neuromodulation.

Second, temporal dynamics of calcium signal changes in multiple cells suggested an average response time of 0.9 ± 0.2 s (Fig. 2e; **Supplementary Table 3**). It has been reported that TRPV1 can be activated by a temperature jump on a millisecond timescale^34^. Therefore, the temporal response of our photothermal TRPV1 activation method should only be limited by the rate of temperature increase, which depends on the power density of the NIR-II illumination. In our cell experiments where an average response time of 0.9 ± 0.2 s was found (Fig. 2e), we used a laser power density of 400 mW mm^−2^, much lower than that of 10^5^-10^6^ mW mm^−2^ in previous reports with ~ms temporal dynamics for activating TRPV1 (**Supplementary Table 3**)^34^. Thus, a longer response time is anticipated for *in vivo* photothermal neuromodulation due to a more stringent constraint on the allowable power density of NIR-II illumination on the animals.

### *In vivo* NIR-II neural stimulation in the mouse hippocampus and motor cortex

Having demonstrated selective activation of MINDS-sensitized TRPV1 with NIR-II illumination *in vitro*, we asked if MINDS allowed for neural activation and behavioral modulation of live mice *in vivo*. A power density of 8 mW mm^−2^ within the safety limit was used for the 1064 nm illumination^40^. To avoid overheating the scalp and the brain with the NIR-II light, a thermal camera was used to monitor the temperature of the scalp during the experiment (**Supplementary Fig. 10**), and a discontinuous illumination protocol with feedback control was applied to activate TRPV1 with a temperature increase from 37 °C to 39 °C (**Methods**)^41^. A temperature increase of 2 °C was sufficient to partially activate TRPV1 expressed in the neuron membranes (**Supplementary Note 1**). In addition, thermal damage to the brain tissue is limited for a temperature of ≤39 °C according to previous reports and guidelines (**Supplementary Note 2** and **Supplementary Table 4**). We have also independently confirmed minimal thermal damage in the M2 region under our stimulation protocol via immunohistochemical studies (**Supplementary Fig. 11**).

We first examined whether NIR-II neuromodulation would be sufficient to activate neural activity *in vivo*. A viral vector with a pan-neuronal promoter eSyn (AAV5-eSyn-TRPV1-p2A-mCherry) was stereotactically injected into the mouse hippocampus for transduction of neurons with TRPV1, followed by the injection of MINDS in the same region 3~4 weeks later (see **Methods**). *In vivo* electrophysiological recording was performed in anesthetized mice immediately after injection of MINDS. The hippocampus was chosen for recording due to the relatively high neuron soma density in CA1 that affords straightforward measurement of extracellular action potentials^42^. Recordings in the TRPV1+/MINDS+ mouse brain showed a statistically significant increase in neuron firing rate upon NIR-II illumination in a reproducible manner, compared to the control groups that lack TRPV1 or MINDS (Fig. 3a and **Supplementary Fig. 12**). This lack of a significant change in firing rate of the TRPV1–/MINDS+ group ruled out the possibility of non-specific neuron modulation directly through the membrane capacitive current or inwardly rectifying K+ channels driven by the temperature increase^39,43^. Additionally, a similar baseline firing rate without NIR-II illumination between TRPV1+ (5.5 Hz) and TRPV1– groups (6.7 Hz) suggest minimal adverse effects of ectopic TRPV1 expression to the neuron’s excitability. Similarly, a comparable baseline firing rate between MINDS+ (5.8 Hz) and MINDS– groups (6.4 Hz) suggest minimal adverse effects to the neuron’s firing ability due to the presence of MINDS.

**Fig. 3.**
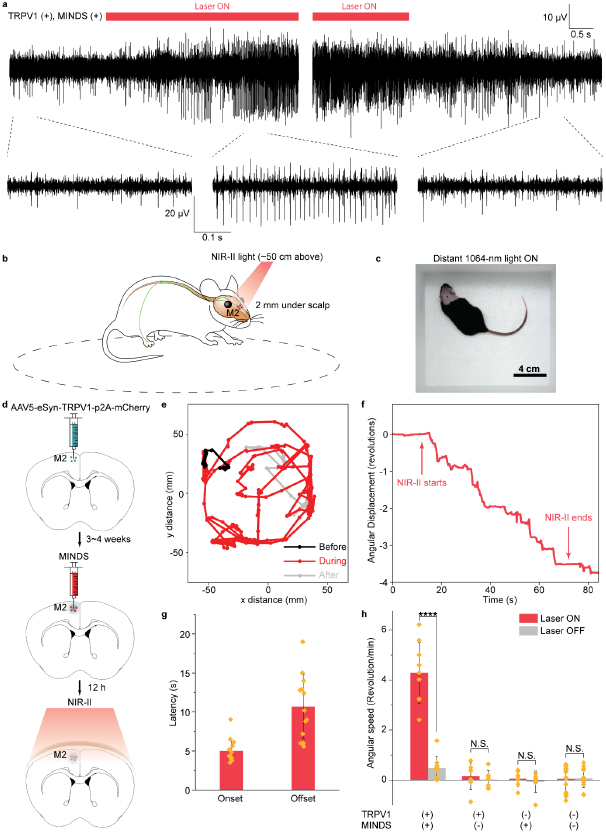
Through-scalp NIR-II neuromodulation of the mouse hippocampus and motor cortex. (**a**) Representative traces showing the increase in neuron firing rate in the mouse hippocampus upon 1064-nm light illlumination (left). The neuron firing rate returned to the baseline after the 1064-nm light was turned off (right). (**b**) Schematic showing through-scalp NIR-II activation of motor neurons (located ca. 2 mm underneath scalp surface) that ectopically overexpress TRPV1 channels and project to the spinal cord for controlling the unilateral limb via the sensitization of MINDS (red dots). A unilateral circling behavior (dashed circle) is evoked for the subject upon NIR-II illumination. (**c**) A representative image showing the arena for motor behavioral modulation by distant NIR-II illumination (invisible in the image since the camera used to take this image is insensitive to the NIR-II wavelength). (**d**) Schematics illustrating the process of TRPV1 virus delivery (green dots, top), injection of MINDS (red dots, middle, where the green haze indicates neural tissue transduced with TRPV1), and NIR-II activation of TRPV1 neurons through the intact scalp (bottom). (**e**) Trajectory of the mouse before (black), during (red) and after (gray) distant NIR-II illumination, showing obvious unilateral circling behavior with the NIR-II light on. (**f**) Angular displacement (number of revolutions, where positive indicates counter-clockwise revolutions and negative clockwise revolutions) before, during and after NIR-II illumination. (**g**) Bar chart showing the statistics of onset and offset latency times for M2 neural stimulation with NIR-II illumination. The onset latency time is defined as the time interval between NIR-II illumination and the start of unilateral circling. The offset latency time is defined as the time interval between the end of NIR-II illumination and the cease of unilateral circling. The error bars indicate SD and each point indicates an independent trial from a total of *N* = 13 trials. (**h**) Statistical analysis of rotational movements of animals under different experimental conditions. Only the animals (*n* = 6) that received both TRPV1 transduction and MINDS injection show a statistically significant increase of unilateral rotation under 1064-nm light (****, *P* < 0.0001). In contrast, all other conditions (*n* = 3) do not show a statistically significant difference between the laser turned on and off (N.S., not significant). Error bars indicate ±1 SD and each point indicates an independent trial.

We then hypothesized that NIR-II illumination was able to activate motor cortex neurons through the intact scalp and induce unilateral circling of freely behaving animals (Fig. 3b). The lack of any brain implants or head tethering, and the use of distant NIR-II light to track the mouse head, should eliminate any perturbation to the natural behavior of freely moving animals within the arena (Fig. 3c and **Supplementary Table 1**). To this end, we performed unilateral transduction of neurons in the secondary motor cortex (M2)^23,44^ using the AAV5-eSyn-TRPV1-p2A-mCherry virus, followed by the injection of MINDS in the same region 3~4 weeks later (Fig. 3d). The synapsin promoter was used in a recent study of magnetothermal deep brain stimulation to drive unilateral circling by stimulating the secondary motor cortex^23^. Distant 1064-nm illumination, invisible to the subject and targeted to the head, was able to penetrate through the scalp and skull for modulating MINDS-sensitized M2 neurons of the mouse without any fiber implants or tethering in a rectangular arena (Fig. 3c; see **Methods**).

Behavioral experiments demonstrated that NIR-II light effectively induced unilateral circling behavior, as evidenced by the contrast between the random exploratory trajectories without NIR-II light and unilateral circling behavior during NIR-II illumination (**Supplementary Movie 1** and Fig. 3e). NIR-II neuromodulation has a key advantage over conventional optogenetics for modulating motor activity, since it frees the animals from any permanent brain implant or head tethering (see **Discussion** for comparison). Furthermore, compared to magnetothermal stimulation where the magnetic coil limits the arena to roughly the size of the animal^23^, distant NIR-II illumination enables a much larger arena for animal behavioral study without any size constraints.

It is important to evaluate the temporal kinetics of *in vivo* NIR-II neuromodulation. From the behavioral study of NIR-II induced unilateral circling, an average latency time of 5.0 ± 1.5 s (mean ± 1 SD; Fig. 3f&g) was found under a power density of 8 mW mm^−2^ at 1064 nm. A similar latency time of ca. 2.9 s, which was defined as the time it took for the firing rate to increase by 50% of the baseline^24^, was found from *in vivo* electrophysiological recordings at the same power density (**Supplementary Fig. 12e**). As discussed above, the temporal resolution of our method depends on the power density of NIR-II illumination. Despite the ~ms response time under a power density of 10^5^-10^6^ mW mm^−2^ in previous reports (**Supplementary Table 3**)^34^ and the response time of 0.9 ± 0.2 s under a power density of 400 mW mm^−2^ in our *in vitro* experiments (Fig. 2e), *in vivo* application of NIR-II illumination should follow strict guidelines on the limits of exposure to the laser to prevent any thermal damage to the brain^40^. Specifically, for NIR-II neuromodulation in the M2, a power density of 8 mW mm^−2^ was used, 50× lower than used *in vitro*, thereby leading to a longer response time. The ~5 s latency time also agrees with temperature measurement in the M2 under NIR-II. Taking advantage of the temperature-dependent fluorescence intensity of DyLight550 (**Supplementary Fig. 13a**)^23,33^, we covalently conjugated MINDS with DyLight550 and performed *in vivo* temperature measurement with a fiber photometry setup in head-fixed animals (see **Methods**). With this method, we found that the M2 temperature increased to 39 °C within 1.5 s (**Supplementary Fig. 13b**). The difference between the temperature measurement and the latency time measured from behavioral experiments could be attributed to the delay between the temperature increase and the ramp of neural activity to reach the threshold, and the delay between neural activation and the onset of behavioral response^45^.

Besides the onset time of *in vivo* NIR-II photothermal activation, we also measured the offset time and compared it with similar methods. Electrophysiological measurements revealed an offset time of 8.6 s, which was defined as the time it took for the firing rate to drop below 50% above the baseline (**Supplementary Fig. 12e**). A similar offset time of 10.7 s was found from behavioral experiments when the NIR-II light was turned off (Fig. 3f). The offset time is shorter than that of magnetothermal stimulation (14.7 s)^23^ as heat diffusion is more spatially confined with a shorter onset time, thus leading to more rapid cooling. Moreover, the offset time of photothermal neuromodulation is much shorter than that of chemogenetics (hours), which is limited by slow clearance and regenerative downstream events of designer drugs (e.g., CNO and varenicline)^25,26^.

We next demonstrated the necessity of ectopically expressed TRPV1 for *in vivo* NIR-II neuromodulation. First, immunohistochemical staining of TRPV1 in the M2 region of the wild-type mouse brain revealed low endogenous expression of TRPV1 (**Supplementary Fig. 14a**). Second, we have carried out *in vivo* electrophysiological, behavioral, and immunohistological studies to compare a group of TRPV1+/MINDS+ animals (*n* = 11) with another group of TRPV1–/MINDS+ animals (*n* = 9). We hypothesized that these two groups of animals would show no statistically significant difference in these studies if the endogenously expressed TRPV1 or other temperature-sensitive ion channels suffice in driving the NIR-II photothermal neuromodulation in the presence of MINDS. The recorded neuron firing rate change upon NIR-II illumination in the TRPV1–/MINDS+ group is significantly smaller than that in the TRPV1+/MINDS+ group (P < 0.001; **Supplementary Fig. 12c**). Furthermore, we have also verified the specific activation of M2 neurons with ectopically expressed TRPV1 in the presence of MINDS by immunohistological staining for c-Fos, an immediate early gene for labeling neuronal activity (**Supplementary Fig. 14**)^24^. A significant increase in the number of c-Fos-positive cells was only observed in the TRPV1-overexpressed mice with MINDS injection after NIR-II illumination, but not in any of the control groups. In addition, mice in the TRPV1+/MINDS+ group exhibited a significant increase in rotation speed (4.60 rev min^−1^) upon NIR-II illumination, which was not observed in the TRPV1–/MINDS+ group (*P* < 0.0001; Fig. 3h). The behavioral results also help rule out the possibility that other TRPV1-expressing neurons outside the brain, such as the dorsal root ganglion neurons of the peripheral nervous system (PNS)^46^, drove the observed unilateral rotation due to the diffusion of brain-injected MINDS to the PNS. Taken together, these results confirmed the selective NIR-II stimulation of ectopically TRPV1-expressing neurons in the M2 instead of those with or without endogenous expression of TRPV1.

We then investigated the necessity of MINDS for *in vivo* NIR-II neuromodulation. First, *in vivo* temperature measurements revealed that the M2 temperature increased from 37 to 39 °C within 1.5 s in the presence of MINDS, whereas it only increased to 37.5 °C after the same illumination duration without MINDS (**Supplementary Fig. 13b&c**). These results indicate the superior photothermal performance of MINDS over normal brain tissue. Second, we carried out *in vivo* electrophysiological, behavioral, and immunohistological studies to test whether direct NIR-II illumination without MINDS can TRPV1 in the brain. We found that the change of neuron firing rates upon NIR-II illumination in the TRPV1+/MINDS– group is significantly smaller than that in the TRPV1+/MINDS+ group (P < 0.001; **Supplementary Fig. 12c**). Additionally, the significant increase in rotation speed of mice in the TRPV1+/MINDS+ group upon NIR-II illumination was not found in the TRPV1+/MINDS– group under the same experimental conditions (*P* < 0.0001; Fig. 3h). Moreover, immunohistological imaging revealed little c-Fos expression in TRPV1+ neurons in the absence of MINDS (**Supplementary Fig. 14**). Therefore, the strong absorption of NIR-II light by MINDS sensitizes TRPV1 to low-intensity NIR-II illumination for neuromodulation. In summary, a significant increase in the average neuron firing rate, mouse rotation speed, and the number of c-Fos-positive cells was only observed in the TRPV1+/MINDS+ mice after NIR-II illumination, but not in any of the control groups (**Supplementary Fig. 12c**, Fig. 3h, and **Supplementary Fig. 14**). Together these data demonstrate that both TRPV1 and MINDS are essential for effective and selective NIR-II photothermal stimulation to modulate animal motor behaviors.

### NIR-II deep-brain stimulation

We next investigated whether NIR-II light can penetrate deep enough to stimulate neural activity in deep-brain regions. We chose the VTA as our target owing to its deep location inside the brain and its pivotal role in the brain’s reward circuitry that could be studied with a conditioned place preference test (Fig. 4a)^2^. It has been demonstrated that optogenetic activation of VTA dopaminergic neurons induces real-time place preference, yet an invasive brain implant or a tethered interface is typically required for light delivery in this deep brain region^2,13,18^. We selectively tagged dopaminergic projections from VTA to the nucleus accumbens (NAc) in the ventral striatum by using an AAV virus (AAV5-EF1α-DIO-TRPV1) encoding TRPV1 in a double-floxed inverted open reading frame (DIO) in tyrosine hydroxylase (TH)-driven Cre recombinase (TH-Cre) transgenic mice (Fig. 4b)^13^. The Cre-dependent specific expression of TRPV1 in VTA dopaminergic neurons was confirmed by the colocalization of TH and TRPV1 (**Supplementary Fig. 15**). After injecting MINDS in the VTA, we performed contextual conditioning for naturally behaving animals in a Y-maze by associating a specific grating pattern at one of the arm terminals with widefield NIR-II illumination (red dashed square, Fig. 4c; **Supplementary Fig. 16**). Due to the deep location of VTA in the brain, a power density of 10 mW mm^−2^, slightly higher than that used in the M2 (8 mW mm^−2^) but still within the safety limit^40^, was applied^29^. Similar to NIR-II stimulation in the M2, we have confirmed minimal thermal damage to the neural tissue above the VTA, which was closer to the NIR-II illumination (**Supplementary Fig. 11**, **Supplementary Fig. 13d**, **Supplementary Table 4** and **Supplementary Note 2**).

**Fig. 4.**
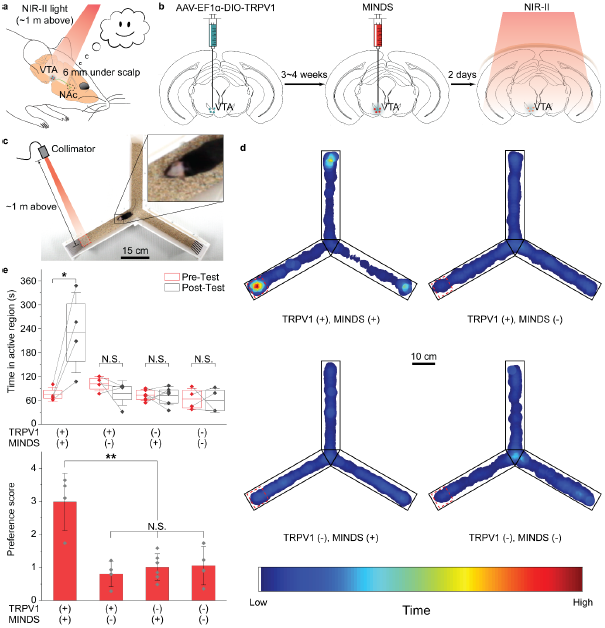
Through-scalp NIR-II stimulation of a deep-brain region. (**a**) Schematic showing through-scalp NIR-II activation of VTA dopaminergic neurons (located ca. 6 mm underneath scalp surface) that ectopically express TRPV1 and project to NAc (green) via the sensitization of MINDS (red dots). (**b**) Schematics illustrating the process of the specific transduction of dopaminergic neurons with TRPV1 (green dots, left), injection of MINDS (red dots, middle, where the green haze indicates neural tissue transduced with TRPV1), and distant NIR-II illumination for neural modulation through the intact scalp (right). (**c**) Photograph showing the setup of a Y-maze for contextual conditioning test. In this setup, the mouse was trained in an implant-free and tether-free manner for three consecutive days with overhead NIR-II light which illuminated from ~1 m above the head in one of the arms (red dashed square) and tested for locational preference on the following day after the training. (**d**) Representative post-test heat maps showing the time of travel of mice under different experimental conditions. Red dashed squares indicate NIR-II irradiated regions. (**e**) Statistical analysis of time spent in the NIR-II illuminated region (top) and preference score (bottom) for different TRPV1/MINDS combinations in the VTA. In the top graph, the center lines, open rectangles, and whiskers indicate the mean, 25/75 percentiles and ±1 SD, respectively, with data points shown for the average value of each animal. Only trials in TRPV1+/MINDS+ animals show a statistically significant increase of time spent in the NIR-II illuminated region in post-test trials vs. pre-test trials (*, *P* < 0.05; N.S., not significant). In the bottom graph, error bars represent ±1 SD, with each data point representing the preference score for each animal. Only the TRPV1+/MINDS+ animals (*n* = 4) show a statistically significant level of place preference compared to the control groups (**, *P* < 0.01). The preference score is defined as the ratio of total time each mouse spent in the NIR-II light illuminated arm terminal during post-test vs. pre-test trials.

After contextual training with NIR-II neuromodulation on three consecutive days^13^, mice with both TRPV1 transduction and MINDS injection in VTA demonstrated strong preference in the NIR-II illuminated arm terminal, as evidenced by longer time the mice spent therein in the post-test (Fig. 4d, **Supplementary Fig. 17** and **Supplementary Movie 2 & 3**). This result proves successful deep-brain neuromodulation in implant-free and tether-free animals with our approach. Furthermore, the specificity of *in vivo* NIR-II neuromodulation of VTA neurons was validated by comparing the TRPV1+/MINDS+ group with the control groups, in which the animals missed TRPV1 transduction, or MINDS injection, or both (Fig. 4e).

We first confirmed the selective NIR-II stimulation of ectopically TRPV1-expressing dopaminergic neurons in the VTA instead of other neurons with or without endogenous expression of TRPV1. Importantly, immunohistochemical staining of TRPV1 in the VTA region of the non-transduced brain revealed low endogenous expression of TRPV1 in the TH+ neurons (**Supplementary Fig. 15a**). Similar to NIR-II neuromodulation in the M2, endogenous TRPV1 expression in other parts of the brain or in the PNS could have led to the observed place preference. However, the demonstrated place preference in the TRPV1+/MINDS+ group was not found in the TRPV1–/MINDS+ group, with a statistically significant difference between the groups (Fig. 4e). Furthermore, we have also verified the specific activation of dopaminergic, TH+ neurons with ectopically expressed TRPV1 by immunohistological staining for c-Fos, which was not found for TH+ neurons lacking TRPV1 overexpression (**Supplementary Fig. 15a&b**). Finally, despite the non-specific heating of the neural tissue, especially the tissue above the VTA due to proximity to the NIR-II light, negligible depolarization of the TRPV1– neuron membrane has been found (**Supplementary Fig. 12**), confirmed in previous magnetothermal neural stimulation reports^23,24^.

We next sought to prove the necessity of MINDS to sensitize the ectopically expressed TRPV1 receptors under NIR-II. First, we have found that without MIINDS, the VTA only reached a temperature increase of 0.2 °C, in contrast to 2.1 °C with MINDS over the same NIR-II illumination condition (**Supplementary Fig. 13d**). Second, the results of behavioral experiments demonstrated negligible place preference in the two MINDS– groups (Fig. 4e). Finally, immunohistological staining of VTA brain slices revealed a significantly lower percentage of c-Fos+ cells out of all TRPV1+ neurons without MINDS (**Supplementary Fig. 15c**). Given the importance of MINDS for NIR-II sensitization, we also evaluated the potential long-term utility of MINDS. We have verified that MINDS remained functionally stable *in vivo* and still demonstrated superior differential heating over normal brain tissue for at least 2 weeks after injection (**Supplementary Fig. 18**), indicating its potential for long-term utility. Taken together, we have demonstrated neural stimulation of deep-brain regions located ca. 6 mm underneath the scalp with distant NIR-II light, which eliminates the needs for any brain implant or head tether, while permitting mouse behavioral study in a large-size arena such as the Y-maze (**Supplementary Table 1**).

## Discussion

Here, we report a novel NIR-II light-based neuromodulation approach for the experimental manipulation of neural activity in a depth range from 1 mm to 6 mm underneath the intact scalp of freely behaving animals. We paired the neuron-specific ectopic expression of heat-sensitive TRPV1 ion channels, which were recently reported to endow NIR vision to rodent and human retinas^28^, with NIR-II absorbing polymeric nanoparticles (MINDS) as light sensitizers to achieve remote control of neuronal activity *in vivo*. In coupling these two systems, we demonstrated controlled activation of TRPV1 channels in neurons, resulting in striking behavioral and electrophysiological changes. Many optogenetic and chemogenetic systems now exist for controlling the neuronal activity, each with its particular advantages and disadvantages. The NIR-II neuromodulation method we report here provides an additional tool with some salient advantages over existing approaches.

### Utility of NIR-II neuromodulation for *in vivo* applications

NIR-II optogenetic neuromodulation sits in a “sweet spot” between conventional optogenetics and chemogenetics.

Conventional optogenetics uses visible light to activate a wide range of opsins, with the longest wavelengths reported for one-photon activation of red-shifted opsins at 635 nm^14,15^. Due to the strong scattering and absorption of visible photons (400-750 nm) in the brain, skull, and scalp^9,30^, a chronic brain implant of an optical fiber^44^ or a microLED^13^ is usually required to deliver light to deep-brain regions, whereas transcranial red-light delivery can penetrate to a depth of 7 mm by mounting the optical fiber above the exposed skull of head-tethered animals^14,15,18,19^. In the latter example, optical fibers need to be fixed on the exposed skull after scalp removal, since only ~0.02% of the incident 635-nm light can penetrate to the 7 mm depth^18^, necessitating a high output power of ≥400 mW mm^−2^ from the fiber. In contrast, 1064-nm light used in this work retains 1.17% of the incident photons at the same depth (**Methods**), thus enabling neuromodulation of untethered animals with a distant light source placed >50 cm above the mouse head with an incident power density of ≤10 mW mm^−2^. Additionally, two-photon activation of opsins enables deeper brain penetration, yet this technique requires a coherent, focused laser beam in head-fixed animals^17^. Moreover, although 980-nm NIR irradiation has been used for deep-brain optogenetics assisted by upconversion nanoparticles, 980-nm light is near one of major absorption bands of water abundant in the brain tissue^9^, thus leading to non-specific tissue heating, a limited penetration depth (**Supplementary Fig. 2**), and a tethered fiber interface for efficient light delivery^21^.

Despite recent advances, two challenges remain for conventional optogenetics: first, the chronic brain implants result in the permanent occupation of the neural tissue and a chronic immune response characterized as gliosis at the implant/tissue interface^11,13^. Second, head fixing and tethering restricts the study of more complex, ethologically relevant behavioral paradigms. For example, head tethering confounds behavioral experiments due to the restriction of the animals’ natural behaviors and limits the study of socially interacting animals (**Supplementary Table 1**). Specifically, behavioral studies that involve animals in a confined space, such as the tube test for studying social hierarchy^12^ and the acute stress test in a restrainer^47^, are typically incompatible with fiber tethering. A fiber-tethered interface is also incompatible with social behavioral studies that involve multiple mice in the same cage (e.g., the IntelliCage) due to potential tangling and biting of the fibers^48^. A similar head-fixed or head-tethered setup is required for sonogenetic^49^ and sono-optogenetic^22^ neuromodulation. In contrast, NIR-II neuromodulation enables free motion and interaction of the subjects via an implant-free and tether-free stimulation interface, since distant 1064-nm light sources can target each animal from ~1 m above. In summary, our approach demonstrates a minimal chronic immune response due to the elimination of brain implants (**Supplementary Fig. 8**) and allows natural behavioral study of freely moving and potentially socially interacting animals due to the removal of head tethering (Fig. 3&4)^12^.

Additionally, our approach enables much shorter onset and offset response times than chemogenetic neuromodulation. Similar to our approach, chemogenetics does not require any brain implants or head tethering, but is limited by the relatively coarse temporal resolution of neuronal manipulation^25,26^. In this work, we report an onset time of ~5 s and an offset time of ~11 s for *in vivo* NIR-II neuromodulation under a safe laser power density of 8 mW mm^−2^ (Fig. 3). These response times are remarkably shorter than those reported for chemogenetics (~min onset and ~h offset)^25,26^. The latency of our approach does not rely on the slow-pharmacokinetics of any molecular actuators but is instead determined by the rate of heating and cooling of NIR-II sensitizers, MINDS, in the brain tissue. The strong absorption of MINDS at the 1064-nm wavelength (Fig. 1e), along with the 2.3-fold increase in TRPV1 channel conductance from 37 °C to 39 °C (**Supplementary Note 1**)^29^, contribute to the seconds-level latency times of neural activation and inactivation. These seconds-level latency times of NIR-II neuromodulation agree with *in vivo* temperature measurements in the brain (**Supplementary Fig. 13b**) and represent a major advantage over chemogenetics.

Compared to magnetothermal neuromodulation^23,24^, our approach allows for manipulating the behaviors of freely moving animals with an unlimited arena size (**Supplementary Table 3**). This advantage is due to negligible attenuation of NIR-II light in free space vs. the 1/*r* decay of the alternating magnetic field with distance from the coil, according to the Biot-Savart law. In addition, the photothermal conversion efficiency of MINDS (3.1×10^4^ W g^−1^ at 8 mW mm^−2^) is higher than the magnetothermal conversion efficiency of superparamagnetic nanoparticles driven by alternating magnetic field (ca. 200 W g^−1^ at 7.5 kA m^−1^ and 1 MHz)^23^. As a result, NIR-II light can be applied to an unlimited size of the behavioral apparatuses, whereas a spatially confined magnetic coil is required to produce a sufficiently strong magnetic field. Another consequence of the higher heating efficiency of our approach is the shorter onset response time of NIR-II photothermal neuromodulation (~5 s) than that of magnetothermal stimulation (~22 s) in mouse behavioral studies^23^.

Compared to other photothermal neural stimulation approaches^5,38,50–52^, our NIR-II photothermal method has the following advantages (**Supplementary Table 3**). First, our approach is the only photothermal neural modulation method that has been demonstrated in free-moving animals, owing to the tissue attenuation minimum at 1064 nm (**Supplementary Fig. 1**) and the unique optical properties of MINDS (Fig. 1e). Most of the existing photothermal neural stimulation reports were limited to *in vitro* due to the use of short-wavelength light in the visible and NIR-I, which exhibit severe absorption and scattering from the brain tissue^5,6,38,50–52^. Much longer infrared wavelengths of 1889 nm and 1869 nm have also been reported to stimulate cells via photothermally induced membrane capacitive current^39^; however, water absorption becomes a major challenge when applying these wavelengths *in vivo*. Our rational choice of 1064-nm light, in contrast, represents the longest wavelength used for *in vivo* neural modulation and offers the greatest brain tissue penetration for the entire visible to NIR spectrum (**Supplementary Fig. 1&2**). Second, our approach uses the lowest power density for both *in vitro* and *in vivo* studies, with the power density used for *in vivo* study within the reported safe exposure limit of 10 mW mm^−2^ at 1064 nm^40^. This low power requirement is due to the efficient sensitization of MINDS, which demonstrates one of the highest photothermal conversion efficiencies in the NIR-II spectrum (**Supplementary Table 2**). The low power density also allowed fine control of the brain temperature without causing thermal damage (**Supplementary Fig. 11**, **Supplementary Table 4**, and **Supplementary Note 2**). Third, selective transduction of TRPV1 enables cell-type specific modulation of neural activity (**Supplementary Fig. 14&15**), in contrast to non-selective photothermal neuromodulation based on optocapacitive mechanisms^5,6,38,39^. Using multiple readouts including animal behaviors, *in vitro* calcium imaging, *in vivo* extracellular recordings, and immunohistological imaging, we have proved the ability of NIR-II neuromodulation to target specific cell types without interference from undesired non-specific activation.

### Potential limitations of *in vivo* NIR-II neuromodulation

While the NIR-II neuromodulation approach reported here overcomes some of the previous challenges in optogenetic, chemogenetic, magnetothermal, and photothermal neuromodulation methods, we acknowledge potential limitations of this system.

The primary disadvantage of the *in vivo* NIR-II neuromodulation method is the slow response time (~s), which is still much longer than most optogenetics approaches (~ms). These relatively slow kinetics of neural activation makes this approach unfavorable for certain neurobiological events that happen on the order of milliseconds^1^. However, some behavioral experiments and many neurological diseases occur on a time scale of days to even years^53^, while neuromodulation techniques with much longer latency time of minutes (e.g. chemogenetics) have also achieved widespread utility for probing neural circuits in animal models^26^. Thus, we believe that with the advantage of a tether-free brain interface, our approach is particularly suitable for long-term neuroscience studies with time frames of days to months, such as behavioral studies of socially interacting animals in the IntelliCage^48^.

Another disadvantage of the *in vivo* NIR-II neuromodulation method, which is closely related to the slow kinetics as discussed above, is the non-specific heating of brain tissue due to thermal diffusion. As discussed earlier, we used untargeted MINDS for activating TRPV1 under NIR-II because the temporal dynamics of TRPV1 activation precludes the need for physically binding MINDS to TRPV1 channels. The thermal diffusion leads to non-specific heating of the brain tissue and has been reported to complicate the interpretation of the behavioral results due to the suppression of neural activity^43^. Nonetheless, by comparing the results of electrophysiological, behavioral, and immunohistological studies between the TRPV1+ and TRPV1– animal groups (Fig. 3h, 4e, **10c, 14 & 15**), we have not found any statistically significant activation or suppression of neural activity solely due to heating of brain tissue to ~39 °C. Besides, heating of the brain tissue to 39 °C, which was sufficient to significantly increase the conductance of TRPV1 channels (**Supplementary Note 1**)^29^, does not cause thermal damage to the brain tissue (**Supplementary Fig. 11**, **Supplementary Table 4**, and **Supplementary Note 2**).

Although we have achieved noninvasive control over specific neural cell types during the behavioral experiments, TRPV1 transgene delivery and MINDS sensitization still involve invasive intracranial injections. Nonetheless, we anticipate that this invasive procedure can be mitigated by systemic delivery of TRPV1 using AAV-PHP.eB virus^54^ or creating a transgenic mouse line with specific expression of TRPV1 in neurons^55^, and delivering MINDS through ultrasound-mediated blood brain barrier (BBB) openings owing to their relatively small sizes^56^. We have demonstrated that MINDS remain functionally stable *in vivo* for at least 2 weeks after delivery into the brain with only a slight decrease in heating capability under NIR-II light, probably due to gradual diffusion from the injection site over time (**Supplementary Fig. 18**). More comprehensive studies are needed in the future to evaluate the functional lifetime and clearance pathways of MINDS from the brain for chronic *in vivo* NIR-II neuromodulation.

To conclude, in this study we describe the validation and utility of deep-brain neuromodulation with distant NIR-II illumination, which penetrates deep into the brain through the intact scalp and skull to selectively activate MINDS-sensitized TRPV1 channels in neurons. The utility of our method sits between optogenetics and chemogenetics: it eliminates the chronic brain implants and head tethering/fixing required for optogenetics and features a more precise temporal control of activation and inactivation than chemogenetics. Therefore, the NIR-II neuromodulation approach reported here allows timely behavioral modulation of freely moving subjects with minimal chronic gliosis in the neural tissue and no interference to natural animal behaviors. With complete elimination of any brain implant and head tethering owing to the deep-brain penetration of NIR-II photons, our approach could afford wide applications in dissecting the complex neural circuits of normally behaving animals in a naturally interacting environment such as the IntelliCage^48^. When combined with a red-shifted bioluminescent reporter for noninvasive neural activity imaging^57^, our method may realize an all-optical bidirectional neural interface through the scalp and skull in behaving animals.

## Acknowledgements

We thank W. T. Newsome, M. Z. Lin, X. Chen, L. Luo, H. Dai, D. Jiang, and J. R. Sanes for helpful discussions. We thank the Stanford Animal Histology Services for help with preparation of histologic specimens. G.H. acknowledges support from the Wu Tsai Neurosciences Institute of Stanford University. X.W. acknowledges support from the Stanford Graduate Fellowship. K.S.O. acknowledges the NeuroTech training program supported by the National Science Foundation under Grant No. 1828993. This work was performed in part at the Stanford Nano Shared Facilities (SNSF) and Cell Sciences Imaging Facility (CSIF) of Stanford University. K.P. thanks Nanyang Technological University (startup grant: M4081627) and Singapore Ministry of Education Academic Research Fund Tier 2 (MOE2016-T2-1-098) for financial support. Some schematics were made in part using BioRender.

## Author contributions

X.W., Y.J., K.P. and G.H. conceived and designed the project, X.W., Y.J., N.J.R., J.L., W.R., S.C., R.Y., A.S. and K.S.O. performed the experiments. X.W., Y.J., N.J.R., J.L., W.R., S.C., K.P. and G.H. analyzed the data and wrote the manuscript. All authors discussed the results and commented on the manuscript.

## Competing interests

Authors declare no competing interests.

## Data and materials availability

The data that support the findings of this study are presented within the manuscript and are available from the corresponding author upon request.

## Code availability

The custom MATLAB code for Monte Carlo and temperature dynamics simulation and data analysis is available from the corresponding author upon request.

## Notes

### Competing Interest Statement

The authors have declared no competing interest.

