## Supplementary Material for "Through-scalp deep-brain stimulation in tether-free, naturally-behaving mice with widefield NIR-II illumination"

**This PDF file includes:**

Methods

Supplementary Notes 1 to 2

Supplementary Figures 1 to 18

Supplementary Tables 1 to 7

Captions for Supplementary Movies 1 to 3

Supplementary References

### Methods

#### Synthesis of *m*acromolecular *i*nfrared *n*anotransducers for *d*eep-brain *s*timulation (MINDS).

Chemicals were purchased from Sigma-Aldrich unless otherwise claimed. Benzobisthiadiazole (CS10361) was purchased from Luminescence Technology Corp (Lumtec, New Taipei City, Taiwan). Briefly, 4,7-bis(5-bromo-4-(2-octyldodecyl)thiophen-2-yl)bisbenzothiadiazole (CS10361, 25 mg, 0.023 mmol), *trans*-1,2-bis(tributylstannyl)ethene (16 μL, 0.03 mmol), tris(dibenzylideneacetone)dipalladium(0) ($Pd_2(dba)_3$, 0.6 mg, 0.00066 mmol), and tri(*o*-tolyl)phosphine (1.8 mg, 0.006 mmol) were weighed and transferred to a 50 mL Schlenk flask followed by dissolution in chlorobenzene (8 mL). After 3 cycles of freeze-pump-thaw degassing, the abovementioned mixture was heated at 100 ℃ in oil bath under the protection of nitrogen atmosphere to perform Stille polycondensation (**Supplementary Fig. 3**). After reaction for 100 minutes, the obtained mixture was added dropwise to cold methanol under vigorous stirring, followed by centrifugation at 9000 rpm for 10 minutes at 0 ℃ to collect dark precipitates[1]. The obtained precipitates were then washed with cold methanol for 3 times and dried under vacuum pump to afford purified poly(benzobisthiadiazole-*alt*-vinylene) (pBBTV) powder. $^1H$ NMR for pBBTV ($CDCl_3$, σ ppm): 7.6-7.0 (s, 4H), 2.6-2.1 (m, 4H), 1.9-1.8 (m, 2H) (**Supplementary Fig. 4**).

To obtain MINDS, the as-prepared pBBTV (0.1 mg) was weighed and co-dissolved with poly(lactide-co-glycolide)-*b*-poly(ethylene glycol) (PLGA-PEG, PLGA $M_n$ 4500, PEG $M_n$ 2000) (4 mg) in tetrahydrofuran (THF, 2 mL). Subsequently, the mixed solution was quickly injected into deionized water (10 mL) under vigorous ultrasonication to afford homogenous suspension, followed by evaporation of THF under a gentle $N_2$ flow for 40 minutes. The remaining aqueous solution was then sufficiently sonicated and filtered through polyvinylidene fluoride syringe driven filter (220 nm, Millipore, Burlington MA) to remove impurities and precipitates. To afford stock solution, the filtered MINDS solution was concentrated via ultracentrifugation at 3500 rpm at 4 ℃ for 25 minutes (Amicon® Ultra-15 Centrifugal Filters, Molecular weight cut-off 50 kDa, Merck KGaA, Darmstadt, Germany). Aliquots of stock solution were subsequently diluted with buffer solution to designated concentrations.

**Transmission electron microscopy (TEM) imaging of MINDS.**

One drop (7 μL) of homogenous MINDS solution (15 μg $mL^{-1}$) was deposited on a formvar/carbon film coated copper grid (Inlab Supplies Pte Ltd, Singapore) and dried in a desiccator for at least 48 h. Afterwards, TEM images of MINDS were captured on a JEOL JEM 1400 transmission microscope (JEOL USA, Inc., Peabody, MA).

**Dynamic light scattering (DLS) of MINDS.**

Aliquot of the abovementioned MINDS solution was diluted with 1× phosphate buffered saline (PBS, pH 7.4) to a final concentration of 5 μg $mL^{-1}$. Then, the hydrodynamic diameter of diluted MINDS was measured by DLS on a Malvern Nano-ZS Particle Sizer (Malvern Panalytical Ltd, Malvern, UK).

**UV-Vis-NIR absorption spectroscopy of MINDS.**

UV-Vis-NIR absorption spectrum of MINDS solution was measured by a Cary 6000i spectrophotometer (Agilent, Santa Clara, CA) with a total path length of 1 mm, background-corrected for contribution from water. The measured range was 300–1800 nm.

**Determination of photothermal conversion efficiency**

The photothermal conversion efficiency of MINDS was calculated following reported methods in the literature[2,3]. In brief, an aqueous solution of MINDS dispersed in 2 mL PBS (optical density at 1064 nm = 1) or a blank PBS solution without MINDS (2 mL) was placed in 3.5-mL quartz cuvette (Sangon Biotech, Shanghai, China) and illuminated by a 1064-nm laser (Shanghai Connet Fiber Optics Co., Ltd., Shanghai, China) at a power density of 10 mW $mm^{-2}$. The average temperature of the solution was monitored by a dual input J/K type thermometer (TM300, Extech Instruments, Waltham, MA) continuously and plotted as a function of time during both the heating and cooling phases. Considering the MINDS solution in the vial as the system to be studied, energy conservation of the system during the heating phase yields

$$\sum_i m_i C_{p,i} \frac{dT}{dt} = \dot{Q}_{MINDS} + \dot{Q}_{water} + \dot{Q}_{vial} - \dot{Q}_{loss}$$

where $m_i$ and $C_{p,i}$ represent the mass and specific heat capacity, respectively, of all components in the system including the MINDS, water and the vial, $T$ is the temperature of the solution, $\dot{Q}_{MINDS}$, $\dot{Q}_{water}$ and $\dot{Q}_{vial}$ are the photothermal energy input of the MINDS, water and the vial per unit time, respectively, and $\dot{Q}_{loss}$ is the energy loss from the system per unit time due to heat transfer from the system to the surrounding environment. $\dot{Q}_{MINDS}$ can be expressed as

$$\dot{Q}_{MINDS} = I(1 - 10^{-Abs})\eta$$

where $I$ is the power of the 1064-nm laser impinging on the sample, $Abs$ is the absorbance of the MINDS at 1064 nm, and $\eta$ is the photothermal conversion efficiency. On the other hand, the energy loss from the system to environment per unit time due to heat transfer is given by

$$\dot{Q}_{loss} = hA(T - T_{env})$$

where $h$ is the heat transfer coefficient, $A$ is the surface area for heat transfer, and $T_{env}$ is the temperature of the surrounding environment. When the heating curve reaches plateau, $\frac{dT}{dt} = 0$, thus

$$\dot{Q}_{MINDS} + \dot{Q}_{water} + \dot{Q}_{vial} = \dot{Q}_{loss}$$

Plugging in the expression of $\dot{Q}_{MINDS}$ and $\dot{Q}_{loss}$ yields

$$I(1 - 10^{-Abs})\eta + \dot{Q}_{water} + \dot{Q}_{vial} = hA(T_{max} - T_{env})$$

Therefore

$$\eta = \frac{hA(T_{max} - T_{env}) - (\dot{Q}_{water} + \dot{Q}_{vial})}{I(1 - 10^{-Abs})}$$

where $T_{max}$ and $T_{env}$ were obtained from the heating curve of the MINDS solution, $I$ was measured with a thermal power meter (S405C, Thorlabs Inc., Newton, NJ), $Abs$ was measured by a UV-Vis-NIR absorption spectrometer (Agilent, Santa Clara, CA).

The same heating curve was also obtained for the blank solution without MINDS, which reached a maximum temperature of $T'_{max}$ under the same laser heating conditions. At equilibrium for the blank solution, we have

$$\dot{Q}_{water} + \dot{Q}_{vial} = \dot{Q}_{loss} = hA(T'_{max} - T_{env})$$

Plugging the updated expression of $\dot{Q}_{water} + \dot{Q}_{vial}$ in the equation for $\eta$ yields

$$\eta = \frac{hA(T_{max} - T_{env}) - hA(T'_{max} - T_{env})}{I(1 - 10^{-Abs})} = \frac{hA(T_{max} - T'_{max})}{I(1 - 10^{-Abs})}$$

Therefore, to obtain the photothermal conversion efficiency $\eta$, the only unknown variables in this equation are *h* and *A*. In order to measure *h* and *A*, consider the cooling phase when the system receives no energy input from the environment, thus $\dot{Q}_{MINDS} = \dot{Q}_{water} = \dot{Q}_{vial} = 0$:

$$\sum_i m_i C_{p,i} \frac{dT}{dt} = -\dot{Q}_{loss} = -hA(T - T_{env})$$

$$\frac{\sum_i m_i C_{p,i}}{hA} \frac{dT}{T - T_{env}} = -dt$$

Integrating both sides yield

$$\frac{\sum_i m_i C_{p,i}}{hA} ln \left(\frac{T - T_{env}}{T_{max} - T_{env}}\right) = -t$$

Therefore, by plotting $ln\left(\frac{T - T_{env}}{T_{max} - T_{env}}\right)$ as a function of time *t* during the cooling phase, one obtains the value *hA* from the slope of linear fitting. Note that $\sum_i m_i C_{p,i}$ was calculated to be 12.928 J/K based on the mass and the specific heat capacity of quartz container and water.

#### Monte Carlo simulation

Two-dimensional Monte Carlo simulations were performed using MATLAB following a similar procedure as previously reported[4]. The simulation sent photon packets with initial energy E into the scalp at normal incidence. Packets were uniformly distributed within 1 cm centered at the MINDS injection site to match the laser spot size used during experiments. A step size of 100 µm was used to bin the 10 mm deep × 21 mm wide simulation space. Between completely random scattering events, photon packets travelled a distance of $d = -\ln(RAND)/\mu_s'$ ($\mu_s'$ : reduced scattering coefficient). The reduced scattering coefficient can be expressed as $\mu_s' = \mu_s(1-g)$, where $\mu_s$ is the scattering coefficient that describes the likelihood of photons undergoing a forward scattering event, and $g$ is the anisotropy factor (close to 0.9 for most tissues). For each travelled distance between two consecutive scattering events, the packet energy decreased by a factor of $\exp(-\mu_a \cdot distance)$ ($\mu_a$: absorption coefficient). The lost energy was assigned to each traversed pixel (determined by binning) as the absorbed energy. When crossing an interface between scalp, skull, or brain tissue, the refraction of the packet was calculated using the refractive index $n$ of different mediums according to Snell's Law. The optical properties of scalp[5-7], skull[8,9] and brain tissue[10-12] were taken from previous reports and are summarized in **Supplementary Table 5**.

**Simulation of temperature dynamics**

The temperature dynamics in the brain tissue was simulated by solving the heat equation below numerically in MATLAB:

$$\frac{dT}{dt} = \frac{k}{\rho C_v} \nabla^2 T + \Delta T$$

where $k$ is the thermal conductivity, $\rho$ is the density and $C_v$ is the specific heat. The thermal properties of different tissues are summarized in **Supplementary Table 6**.[13] $\Delta T$ is the instantaneous temperature increase rate (units of K/s) induced by NIR-II illumination, which was calculated by converting the absorbed energy resulted from Monte Carlo simulation to a temperature increase according to the appropriate density, specific heat, and photothermal conversion efficiency for each pixel in the scalp, skull, and brain tissue with/without MINDS.

**Estimation of penetration efficiency.**

When travelling inside biological tissues, light is attenuated due to absorption by the tissue (related to the absorption coefficient, $\mu_a$) and isotropic scattering (related to the reduced scattering coefficient, $\mu_s'$). The extinction coefficient ($\mu_e$) of biological tissue is a sum of $\mu_a$ and $\mu_s'$, leading to the through-scalp penetration efficiency ($\eta$) of light at different wavelengths that can be estimated as follows:

$$\eta = \exp\left(-t_{scalp} \cdot \mu_{e,scalp}\right) \cdot \exp\left(-t_{skull} \cdot \mu_{e,skull}\right) \cdot \exp\left(-d_{brain} \cdot \mu_{e,brain}\right)$$

where $t_{scalp} = 0.7$ mm and $t_{skull} = 0.6$ mm are the thickness of scalp and skull, respectively, while $d_{brain}$ is the depth of target brain region below brain surface. Thus, at a depth of 5 mm in the brain, the penetration efficiency values of 635-nm, 980-nm and 1064-nm light were calculated to be 0.064%, 1.4% and 2.9%, respectively. Similarly, the through-skull penetration efficiencies of 1064-nm light at a depth of 7 mm in the brain was calculated to be 1.17% by taking $t_{scalp} = 0$.

**Viral vector construction.**

Viral vectors used in this work include: pAAV-CMV-TRPV1-mCherry plasmid was constructed by Vector Biolabs (Malvern, PA), AAV5-eSyn-TRPV1-p2A-mCherry virus, and AAV5-EF1a-DIO-TRPV1 virus were constructed and packaged by Vector Biolabs (Malvern, PA). For transduction in the secondary motor cortex (M2) with AAV5-eSyn-

TRPV1-p2A-mCherry, it has been reported that the synapsin (Syn) promoter is neuron-specific with little glial expression[14].

**Cytotoxicity test of MINDS.**

Cytotoxicity of MINDS was tested on rat hippocampal primary neurons isolated from Sprague Dawley rat embryos and placed in a 96-well plate. 2 weeks after isolation, the neurons were incubated with 200 μL MINDS solution of different concentrations (0 – 3.6 mg $mL^{-1}$) in neuron culture medium for 24 h, and an additional group of neurons were incubated with 5% DMSO as the positive control. After the incubation, the supernatant was removed, and cells were gently washed with fresh 1× PBS. Colorimetric MTT (3-(4,5-dimethylthiazol-2-yl)-2,5-diphenyltetrazolium bromide) assays (Invitrogen, Carlsbad, CA) were performed according to the manufacturer's instruction. Absorbance at 590 nm was subsequently recorded on a SPECTRAFluor Plus microplate reader (Tecan Group Ltd., Männedorf, Switzerland). Cell viability was calculated as the ratio of absorbance from MINDS-treated or DMSO-treated cells to that of negative control cells incubated in blank medium.

**Preparation of MINDS-DyLight550 conjugates.**

To utilize the temperature-dependent fluorescence intensity of DyLight550 as a molecular temperature probe, we covalently functionalized MINDS with DyLight550. 1 mg of DyLight550 NHS ester (Thermo Fisher scientific, Waltham, MA) was dissolved in 100 μL dimethylformamide (DMF) to make a well-dispersed solution. Then 16 μL of DyLight550 NHS ester in DMF solution was added to 200 μL 1.8 mg $mL^{-1}$ MINDS solution in PBS to achieve a molar-fold excess of $10^4$. The mixed solution was then stored in dark at room temperature for 24 hours. After the incubation, the excess DyLight550 was removed by adding 400 μL DI water followed by centrifugation at 10000 rpm for 10 min (mySPIN 12, Thermo Fisher scientific, Waltham, MA) using a centrifugal filter device with a molecular weight cutoff of 100 kDa. The washing step was repeated for 5 times, and the removal of the excess DyLight550 was confirmed by the lack of DyLight550 absorption peak from the filtrate in the last washing step.

**Cell culture and transfection.**

Human embryonic kidney (HEK) 293T cells were purchased from MilliporeSigma (Burlington, MA). Cell culture was maintained in Dulbecco's modified Eagle's medium (DMEM; Gibco) supplemented with 10% fetal bovine serum (FBS; Gibco). HEK293T cells were transfected with 7.5 μL of lipofectamine ® 3000 (Invitrogen, Carlsbad, CA) with 2500 ng of total DNA of the pAAV-CMV-TRPV1-mCherry plasmid in Opti-MEM medium (Gibco), and used for calcium imaging during near-infrared-II (NIR-II) photothermal stimulation 3-5 days after transfection.

***In vitro* NIR-II stimulation and calcium imaging.**

HEK293T cells were seeded into Confocal Dishes (VWR, Radnor, PA) and allowed to adhere for 24 h in a humidified atmosphere containing 5% $CO_2$ and 95% air at 37 °C. The cells were treated with MINDS at a final concentration of 70 μg $mL^{-1}$ for 24 h in culture medium. Cells were washed 2 times with 1× PBS, stained in Hank's Buffer with 20 mM Hepes solution (HHBS; AAT Bioquest, Sunnyvale, CA; containing 140 mg $L^{-1}$

$CaCl_2$) containing the Cal-520® AM fluorescent indicator (AAT Bioquest, Sunnyvale, CA) prior to live-cell imaging. LSM 780 confocal microscope (Carl Zeiss Microscopy GmbH, Jena, Germany) was used for calcium imaging with an excitation wavelength of 488 nm and an emission window of 500-520 nm for image formation. For NIR-II photothermal stimulation, a built-in 1040 nm laser was focused to a spot size of 250 μm and a power density of 400 mW $mm^{-2}$. The average power density of 1040-nm laser used for photothermal stimulation was significantly lower than previously reported power density using 780 nm ($8\times10^3$ mW $mm^{-2}$) and 808 nm ($10^5$ mW $mm^{-2}$) laser[2,15], owing to the much higher photothermal conversion efficiency of 71% measured for MINDS. Real-time calcium imaging was timed with NIR-II photothermal stimulation from the 1040-nm laser, which was turned on and off on selected regions of the cell culture. Average fluorescence intensity of Cal-520® AM indicator was analyzed using custom-written MATLAB code with the built-in *roipolyarray* function[16].

**Vertebrate animal subjects.**

Adult (20-30 g) C57BL/6J mice (6 weeks old, Charles River Laboratories, Wilmington, MA) and TH-Cre transgenic mice (6 weeks old, Jackson Laboratory, Bar Harbor, ME) were the vertebrate animal subjects used in this study. All procedures performed on the mice were approved by Stanford University's Administrative Panel on Laboratory Animal Care (APLAC). The animal care and use programs at Stanford University meet the requirements of all federal and state regulations governing the humane care and use of laboratory animals, including the USDA Animal Welfare Act, and PHS Policy on Humane Care and Use of Laboratory Animals. The laboratory animal care program at Stanford is accredited by the Association for the Assessment and Accreditation of Laboratory Animal Care (AAALAC International). Animals were group-housed on a 12 h: 12 h light: dark cycle in the Stanford University's Veterinary Service Center (VSC) and fed with food and water *ad libitum* as appropriate.

***In vivo* stereotaxic injection of virus.**

*In vivo* injection of virus into the brains of live mice was performed using a controlled stereotaxic injection method[17]. First, all metal tools in direct contact with the surgical subject were autoclaved for 1 h before use, and all plastic tools in direct contact with the surgical subjects were sterilized with 70% ethanol and rinsed with sterile DI water and sterile 1× PBS before use. Viral vectors were dispersed in sterile 1× PBS with 5% glycerol before injection. Mice were anesthetized by intraperitoneal injection of a mixture of 80 mg $kg^{-1}$ of ketamine (KetaVed®, Vedco, Inc., St. Joseph, MO) and 1 mg/kg dexdomitor (Dexmedesed™, Dechra Veterinary Products, Overland Park, KS). The degree of anesthesia was verified via the toe pinch method before the surgery started. Buprenorphine SR-LAB (Zoopharm, Inc., Windsor, CO) analgesia was given subcutaneously at a dose of 1 mg $kg^{-1}$ body weight before the surgery. To maintain the body temperature and prevent hypothermia of the surgical subject, a homeothermic blanket (Harvard Apparatus, Holliston, MA) was set to 37 °C and placed underneath the anesthetized mouse, which was placed in the stereotaxic frame (World Precision Instruments, Inc., Sarasota, FL) equipped with two ear bars and one nose clamp that fix the mouse head in position. Vet ointment (Puralube®, Dechra Veterinary Products, Overland Park, KS) was applied on both eyes of the mouse to moisturize the eye surface throughout the surgery. Hair removal lotion (Nair®,

Church & Dwight, Ewing, NJ) was used for depilation of the mouse head and iodophor was applied to sterilize the depilated scalp skin. A 1-mm longitudinal incision was made with a sterile scalpel, followed by elongation of the incision by surgical scissors to expose the cranial bone immediately above the targeted brain regions for injection without removing any scalp skin. The incisions were made near the distal periphery of the scalp opposite the injection site for unilateral stereotaxic injections later, thus ensuring the wound in the scalp would not block the underlying neural tissue from overhead NIR-II illumination for behavioral modulation.

The following stereotaxic coordinates were used for injection of viral vectors and MINDS:

Secondary motor cortex (M2)[18-20]: Anteroposterior (AP) +1.0 mm, mediolateral (ML) ±0.5 mm (+0.5 for right hemisphere and –0.5 for left hemisphere), dorsoventral (DV) -0.5 mm

Hippocampus (HIP)[21]: AP -1.5 mm, ML ±1.5 mm, DV -1.5 mm

Ventral tegmental area (VTA)[22,23]: AP -3.5 mm, ML ±0.4 mm, DV -4.2 mm

For each animal, according to the targeted brain region, a 0.5 mm diameter burr hole was drilled using a dental drill (Marathon-III) according to the AP and ML coordinates described above. 2.5 μL of AAV solution was injected into M2, HIP or VTA region of the brain using NanoJet (Chemyx, Inc., Stafford, TX) with a 33-gauge needle (Hamilton Company, Inc., Reno, NV) facing the ventrolateral side at 0.1 μL $min^{-1}$. AAV5-eSyn-TRPV1-p2A-mCherry virus ($2.2 \times 10^{12}$ GC $ml^{-1}$) was used for M2 or HIP transduction in C57BL/6J mice, while AAV5-EF1a-DIO-TRPV1 virus ($3.6 \times 10^{12}$ GC $ml^{-1}$) was used for VTA transduction in TH-Cre transgenic mice. After injection of viral vectors, the syringe needle was allowed to stay inside the brain for 2 min before withdrawal. The incised scalp skin was placed back to cover all exposed cranial bone, and sealed by using Vetbond tissue adhesive (3M, Maplewood, MN). Antibiotic ointment (Johnson & Johnson Consumer, Inc., Skillman, NJ) was applied copiously around the wound. The mouse was returned to the cage equipped with a 37°C heating pad and its activity monitored every hour until fully recovered from anesthesia (i.e., exhibiting sternal recumbency and purposeful movement).

***In vivo* stereotaxic injection of MINDS.**

*In vivo* injection of MINDS into the brains of live mice was performed using a controlled stereotaxic injection method similar to virus injection described above. MINDS were dispersed in sterile Hank's Buffer with 20 mM Hepes solution (HHBS; AAT Bioquest, Sunnyvale, CA; containing 140 mg $L^{-1}$ anhydrous $CaCl_2$) before injection. Mice were anesthetized as previously described, and scalp incision and craniotomy were performed at the corresponding coordinates for M2, HIP or VTA. 2.5 μL of MINDS solution (1.8 mg $mL^{-1}$) was injected into targeted brain region using NanoJet (Chemyx, Inc., Stafford, TX) with a 33-gauge needle (Hamilton Company, Inc., Reno, NV) facing the ventrolateral side at 0.1 μL $min^{-1}$. The same amount of HHBS instead of MINDS was injected for MINDS (-) controls. After injection of MINDS, the syringe needle was allowed to stay inside the brain for 2 min before withdrawal. The incised scalp skin was sealed as described above.

***In vivo* electrophysiological recordings during NIR-II neuromodulation.**

Tungsten microwire electrodes (TM53A15, World Precision Instruments, Sarasota, FL) were used for *in vivo* electrophysiological recording immediately after MINDS or carrier-only injection in anesthetized mice. A 0-80 set screw (18-8 Stainless Steel Cup Point Set Screw; outer diameter: 0.060" or 1.52 mm, groove diameter: 0.045" or 1.14 mm, length: 3/16" or 4.76 mm; McMaster-Carr Supply Company, Elmhurst, IL) was implanted into the following stereotaxic coordinates as the grounding and reference electrode: AP: -4.96 mm, ML: 3.10 mm. After implantation of the grounding screw, two tungsten electrodes were slowly inserted into the left and right hippocampal regions (AP: -1.5 mm, ML: ± 1.5 mm, DV: -1.5 mm). After reaching the targeted regions, the electrodes were glued to the skull using METABOND® dental cement (Parkell Inc., Edgewood, NY). The tungsten probes were connected to an Intan RHD 2132 amplifier evaluation system (Intan Technologies, Los Angeles, CA), and the 0-80 set screw was used as a reference. For photothermal stimulation of the hippocampus, a 1064-nm laser was illuminated in the continuous-wave (CW) mode from a distant collimator, with a power density of 8 mW $mm^{-2}$. Electrophysiological recordings were acquired with a 20 kHz sampling rate and a 60 Hz notch filter. The signals were then analyzed using MATLAB with a bandpass filter of 250-6000 Hz[17]. A total of 52 traces collected from 9 animals were used for statistical analysis.

**NIR-II neurostimulation in the M2.**

One experimental group of mice receiving both virus and MINDS injections in the M2 (*n* = 6) and three control groups of mice (virus injection only, *n* = 3; MINDS injection only, *n* = 3; carrier injection only, *n* = 3) were used in this study. The animals were allowed to completely recover from the stereotaxic injection for 12 h and showed normal behaviors before the behavioral experiments started. It has been reported that a 12-h post-operative period is sufficient for studying the behavioral changes due to neuromodulation with minimal interference from anesthetics[20]. The hair over the scalp and neck was shaven to expose the skin, and a red or blue dot was marked on the skin between the head and the neck for tracking the animal's trajectory. For each session, one mouse was placed in a 14 cm × 15 cm Styrofoam box, and was allowed to explore the space in the box freely. A video camera (Rebel T6, Canon U.S.A., Inc, Melville, NY) was used to record the video (640 × 480 pixels, 25 fps) of the mouse before, during and after distant NIR-II photothermal stimulation. A fiber-coupled collimator connected to the 1064-nm laser (SOL, RPMC Laser Inc., O'Fallon, MO) illuminated a 1 $cm^2$ area that followed the head of the mouse in the Styrofoam box. To demonstrate proof-of-concept, the illumination angle of the fiber-coupled collimator was adjusted manually to track the head of the mouse, while a custom-written MATLAB code that tracks the head position of the animal (described below) could be incorporated into a closed-loop controller for automatic tracking. Precautions were taken during each session to ensure there was minimum change of shadows caused by the camera in the active area of the animal to avoid influence on the animal's behavior. During NIR-II photothermal stimulation, the 1064-nm laser was illuminated in the continuous-wave (CW) mode in free space from the collimator, with a maximum power density of 8 mW $mm^{-2}$. The maximum power density was below the safe exposure limit of 10 mW $mm^{-2}$ at 1064 nm determined by the International Commission on Non-ionizing Radiation Protection[24]. During the unilateral circling test, a thermal camera (FLIR A325sc, FLIR Systems, Inc., Wilsonville, OR) was used to monitor heating in real time by following a previously reported protocol[25] to avoid the thermal damage to the brain tissue (**Supplementary Fig. 10**). Specifically, a closed-loop feedback control was used to turn the laser power down to 0 mW $mm^{-2}$ whenever the temperature approached 39 ºC and turned back up to the maximum allowed power density of 8 mW $mm^{-2}$ when the temperature dropped.

The videos acquired from recording sessions were processed in a custom-written MATLAB code that can track both the red/blue marker on the skin and the center of mass of the animal. Based on the trajectory of the mouse obtained thereof, and the frame rate of the video that defines the time interval between two neighboring locations in the trajectory, the angular displacement of the animal with respect to $t = 0$ s of each session was calculated and plotted. Clockwise rotation was defined as negative angular displacement while counter-clockwise rotation was defined as positive angular displacement. Based on the data of angular displacement as a function of time in each session, angular speed of the animal was computed by dividing the number of revolutions over total length of time when the 1064-nm laser was on. Analysis of the variance (ANOVA) was performed using the built-in function 'anova1' in MATLAB to evaluate the statistical significance in angular speed when the laser was on between experimental and control groups.

**NIR-II neurostimulation in the VTA.**

One experimental group of mice receiving both virus and MINDS injections in the VTA ($n = 4$) and three control groups of mice (virus injection only, $n = 4$; MINDS injection only, $n = 6$; carrier injection only, $n = 4$) were used in this study. The animals were allowed to recover from the stereotaxic injection for 48 h before the behavioral experiments. The hair over the scalp and neck was shaved to expose the skin. For each session, one mouse was placed in a three-compartment Y maze (Maze Engineer, Inc., Cambridge, MA). Each arm of the Y Maze was 35 cm by 5 cm, connected by an equilateral triangle with 5 cm side in the center, resulting in a total area of 536 $cm^2$ for the Y maze. At the end of each arm, a unique context (white, horizontal black stripes, or vertical black stripes) was outfitted to facilitate the formation of contextual visual memory (**Supplementary Fig. 16**)[26].

One complete set of training and probe trials comprises five days: On Day 1, the animal was allowed to explore the space in the Y maze freely for 30 min without any NIR-II illumination. On Day 2 to 4 for training trials, the 1064-nm laser was directed via a fiber collimator to illuminate a 1 $cm^2$ area that followed the head of the mouse whenever its head entered a 25 $cm^2$ area at the end of one of the three arms. Similar to the behavioral experiments in the M2, the illumination angle of the fiber-coupled collimator was adjusted manually to track the head of the mouse, while a custom-written MATLAB code that tracks the head position of the animal (described below) could be incorporated into a closed-loop controller for automatic tracking. The laser was modulated at 20 Hz with peak power density of 33 mW $mm^{-2}$ in the illuminated area and a 30% duty cycle, resulting in an average power density of 10 mW $mm^{-2}$, within the safe exposure limit at 1064 nm determined by the International Commission on Non-ionizing Radiation Protection[24]. Similar to the unilateral circling test described above, a thermal camera was used to monitor the temperature on the scalp of the animal during the conditioning with feedback control of the laser output to avoid thermal damage to the brain tissue. The animal was allowed to freely explore all three arms, including the arm with 1064-nm laser illumination for a total of 30 min on each day. On Day 5 for probe trial, the animal was allowed to explore the space in the Y maze freely for 30 min without any NIR-II illumination (**Supplementary Fig. 17**). During each session, a video camera (Rebel T6, Canon U.S.A., Inc, Melville, NY) was used to record the video (1920 × 1080 pixels, 25 fps) of the mouse. Precautions were taken during each session to ensure there was minimum change of shadows in the active area of the animal to avoid influence on the animal's behavior.

The videos acquired from the probe trial sessions were processed in a custom-written MATLAB code that can track both the red/blue marker on the skin and the center of mass of the animal. Based on the trajectory of the mouse obtained thereof, and the frame rate of the video, the amount of time that the animal spent in each location of the Y maze during the probe trial was plotted in a heat map for visualization of place preference. Preference score was defined as the ratio of time the mouse spent in the NIR-II illuminated arm terminal during post-test vs. pre-test trial. Analysis of the variance (ANOVA) was performed using the built-in function 'anova1' in MATLAB to evaluate the statistical significance in preference between experimental and control groups.

**Temperature measurement during through-scalp NIR-II neuromodulation.**

To demonstrate that the maximum temperature in the brain is below the safety limit, we have performed *in vivo* temperature measurements at different brain regions using a fiber photometry setup, which measures the temperature-dependent fluorescence change of DyLight550 (**Supplementary Fig. 13a**)[20,27]. Following the injection of MINDS-DyLight550 conjugates (or DyLight550 only for MINDS(-) controls) in the M2 or VTA region according to '***In vivo* stereotaxic injection of MINDS**', a fiber optic cannula (CFMXD10, Thorlabs Inc., Newton, NJ) connected to an optical fiber (M89L01, Thorlabs Inc., Newton, NJ) was mounted on the stereotaxic frame (World Precision Instruments, Inc., Sarasota, FL) and inserted into the same brain region. The inserted optical fiber served as the light path for both excitation and emission of DyLight550. A homemade fiber photometric system was used to collect the temperature-dependent fluorescence from DyLight550 upon 1064-nm illumination. In brief, a 532 nm laser (20 μW) was used to excite DyLight550 fluorescence, which was detected by a photomultiplier tube (PMT1001, Thorlabs Inc., Newton, NJ). The optical signals were digitized by a data acquisition module (NI USB-6221, National Instruments, Austin, TX) and simultaneously recorded using a custom-made LabVIEW program (National Instruments) at 60 kHz with a moving average of 15,000 to reduce the noise of measurement. A thermal camera was used to monitor the temperature of the scalp during 1064-nm illumination, while the temperature in the brain was readout from the fiber photometry setup in a synchronized manner to correlate the temperatures in different brain regions with that on the scalp. The same 1064-nm illumination protocol used for stimulating M2 or VTA in the behavioral studies was applied.

**Histology.**

C57BL/6J mice were anesthetized and transcardially perfused with 1× PBS followed by 4% paraformaldehyde (PFA) in PBS. For the evaluation of the chronic immune response (**Supplementary Fig. 8**), transcardial perfusion was performed at 1 week and 3 weeks after the animals received MINDS or carrier injection. For the evaluation of thermal damage in the brain (**Supplementary Fig. 11**), transcardial perfusion was performed at 1 week after the animals received NIR-II illumination. For imaging c-Fos in the M2 and VTA (**Supplementary Fig. 14&15**), transcardial perfusion was performed at 90 min after the animals of different experimental and control groups received NIR-II illumination. Brains were dissected, post-fixed for 24 h at 4 ºC in 4% PFA and then equilibrated in 30% sucrose at 4 ºC for cryoprotection. 20 μm-thick horizontal sections were collected on a cryostat

(Leica CM 3050S, Leica Biosystems Inc., Buffalo Grove, IL). The brain slices were rinsed in 1× PBS three times for 10 min each and blocked using a blocking solution consisting of 0.3% Triton X-100 and 5% goat/donkey serum (from the same species in which secondary antibody is made, Jackson ImmunoResearch Laboratories, Inc., West Grove, PA) in 1× PBS for 1 h at room temperature. If mouse was used as a host species for the primary antibody, an additional 1 h blocking using a 26 μg/ml solution of AffiniPure Fab Fragment Goat Anti-Mouse IgG (Jackson ImmunoResearch Laboratories, Inc., West Grove, PA) was performed. Slices were then incubated with the primary antibodies (**Supplementary Table 7**) containing 0.3% Triton X-100 and 5% goat/donkey serum overnight at 4 ºC. After incubation, slices were rinsed in 1× PBS three times for 10 min each, before they were incubated with the secondary antibodies (**Supplementary Table 7**) containing 0.1% Triton X-100 and 5% goat/donkey serum for 1 h at room temperature. Slices were then rinsed in 1× PBS three times for 10 min each before they were mounted on glass slides with coverslips using ProLong Gold Antifade Mountant (Invitrogen, Carlsbad, CA). The slides remained in the dark at room temperature for at least 24 h before microscopic imaging. Confocal fluorescence images were acquired on a Zeiss LSM 780 confocal microscope.

**Assessment of the chronic functional stability of MINDS *in vivo***

2.5 μL of 1.8 mg $mL^{-1}$ MINDS solution was injected into VTA of C57BL/6J mice as described above. The animals were sacrificed 1 day or 2 weeks after the injection, and 2-mm thick acute coronal sections containing the injected MINDS near the VTA were collected from the freshly dissected brains using a brain slicer (Braintree Scientific Inc., Braintree, MA) and immersed in Hank's balanced salt solution (HBSS) immediately afterwards to prevent dehydration until use. The coronal sections were then transferred to the top of a 10-mm petri-dish lid with the MINDS-injected side facing up. 10 mW mm$^{-2}$ 1064-nm illumination was applied to the brain slice, and a thermal camera was used to monitor the temperature change. Also see the schematics in **Supplementary Fig. 18a** for more information.

**Replication.**

The calcium fluorescence imaging experiments to investigate percentage of responsive cells (with and without MINDS, and with and without NIR-II illumination) have been repeated on 3 independent trials for each experimental condition, resulting in 12 trials in total. The dynamic calcium imaging experiments to investigate the fluorescence intensity change and latency time upon NIR-II illumination have been repeated on 15 cells for each experimental condition, resulting in 60 cells in total. The assessment of chronic immune response has been repeated on 3 mice at each time points and experiment conditions, resulting in 12 mice. The assessment of thermal damage has been repeated on 3 mice for M2 and VTA with or without NIR-II illumination, resulting in 12 mice. The unilateral circling experiments to investigate widefield NIR-II modulation of motor behavior (angular speed and latency time) have been repeated on a total of 15 animals and 37 trials. The *in vivo* electrophysiological studies have been repeated on 9 animals and 52 traces. The *in vivo* temperature measurements have been repeated for 4 times in each group (4 subjects). The assessment of chronic functional stability of MINDS has been repeated on 3 mice for each time point with or without MINDS injection, resulting in a total of 9 mice. In addition, investigations of NIR-II modulation of reward circuitry and place preference have been repeated on a total of 18 independent animals. In total, experiments and analyses of 79 mice were included for replication.

**Statistical analysis.**

The variance in calcium fluorescence intensity (**Fig. 2**), angular speed (**Fig. 3**) place preference (**Fig. 4**), cell density and population percentage (**Supplementary Fig. 11, 14&15**), normalized neural firing rate (**Supplementary Fig. 12**), *in vivo* and *ex vivo* temperature measurements (**Supplementary Fig. 13&18**) for each cell, animal, behavioral trial or electrophysiological traces was calculated, by which the pooled standard deviation (SD) among each experimental group were determined. Comparisons between experimental groups were made using one-way analysis of variance (ANOVA) without normality assumption given its reasonable tolerance of violations to normal distribution[28], and P values of less than 0.05 were considered statistically significant.

#### Supplementary Note 1

##### Estimation of TRPV1 conductance change from 37 °C to 39 °C

The total conductance of TRPV1 over a whole cell is determined by the number of TRPV1 channels *N*, the unitary conductance $i$, and the probability of channel opening $P_0$:

$$G = NiP_0$$

where *N* is independent of temperature *T* but $i$ and $P_0$ are dependent on *T*. On one hand, it has been reported that the unitary conductance of TRPV1 is only weakly dependent on temperature with a $Q_{10}$ temperature coefficient of 1.4 ± 0.1.[29] The probability of channel opening $P_0$, on the other hand, exhibits a sigmoidal dependance on temperature but can be modeled by the Arrhenius equation for the rising phase.[30] Therefore, the total conductance of TRPV1 over a whole cell can be modeled by the Arrhenius relationship for the rising phase.

According to a previous report, at the resting membrane potential of neurons (-70 mV), the conductance of TPRV1 at 25, 30, 35 and 40 °C is 0.0980, 0.188, 0.530 and 4.04 nS, respectively.[29] By fitting these four known data points to an Arrhenius relationship, we found that at -70 mV, the conductance of TRPV1 at 37 °C and 39 °C was 1.14 and 2.62 nS, respectively. Thus, a temperature increase from 37 °C to 39 °C results in a 2.3-fold increase in the total TRPV1 conductance over a whole cell.

The reversal potential of TRPV1 has been reported as -15 mV,[31] higher than the neuron membrane's resting potential of -70 mV. Therefore, when the temperature increases from 37 °C to 39 °C, a 2.3-fold increase of depolarizing current through TRPV1 channels is expected. This 2.3-fold increase in transmembrane current is higher than the reported current changes of designer receptors exclusively activated by designer drugs (DREADDs) when chemogenetically activated to drive neuron responses and behavioral changes in the animals.[32,33] Therefore, a temperature increase of 2 °C is sufficient to modulate the activity of TRPV1-expressing neurons. This finding is also confirmed by a previous magnetothermal neuromodulation study that reported a significant increase of firing rate of TRPV1-expressing hippocampal neurons upon a temperature change from 37 °C to 39 °C.[20]

### Supplementary Note 2

#### NIR-II photothermal neuromodulation without thermal damage

Hyperthermia effect on cellular health depends on both the temperature elevation and the exposure time. We show that there is no thermal damage or pain induced to the scalp or brain tissue under our *in vivo* NIR-II photothermal neuromodulation protocol.

First, the reported safe exposure limit for light impinging on the scalp skin is 10 mW $mm^{-2}$ for exposure duration between 10 s and 30000 s at 1064 nm[24]. In our experiments, the 1064-nm illumination was applied with real-time feedback-based power control to prevent thermal damage within a duration of 300 s, during which the maximum power density was 10 mW $mm^{-2}$. Therefore, the power density of 1064-nm illumination is within the reported safe exposure limit and thus does not cause thermal injury to the scalp.

Second, NIR illumination of the skin could possibly evoke pain and thus alter the animal behavior, and the thermal threshold for pain is much lower than that for grossly detectable physical injury[34]. Similar to thermal damage, the isoeffect time to induce pain in the skin follows an Arrhenius relationship at a given temperature. We used a thermal camera to monitor the mouse scalp temperature during the behavioral study (**Supplementary Fig. 10**), and we found that the exposure times of all behavioral experiments were well below the isoeffect time to induce non-burn pain at the scalp temperatures shown in the thermal images.

Third, besides the scalp, the temperature increase in the brain did not lead to permanent tissue damage in our experiments. We performed immunohistochemical studies on the cerebral cortex, which experienced the highest temperature increase under our stimulation protocol (**Supplementary Fig. 13**). We found no statistical significance in the density of cells labeled by cleaved Caspase-3, Iba1, NeuN or GFAP for mouse brains with vs. without NIR-II illumination, both in the presence of MINDS (**Supplementary Fig. 11**). These results suggested that the stimulation protocols used in our experiments did not lead to increased apoptosis or alter the distribution of microglia, neurons or astrocytes. In addition, it has been reported previously that a temperature increase to 42˚C for 25 min is needed to induce thermal damage to the brain tissue (**Supplementary Table 4**). Furthermore, previous magnetothermal neural modulation studies with similar or even higher temperature increase than our method reported minimal thermal damage to the brain tissue (**Supplementary Table 4**). Our stimulation protocol with the feedback control from the thermal camera ensures no thermal damage to the brain tissue during neuromodulation. Nevertheless, the NIR-II stimulation protocol is expected to be further optimized with a more comprehensive study in the future to understand the physiology of TRPV1 and the interaction between NIR-II light with the brain tissue.

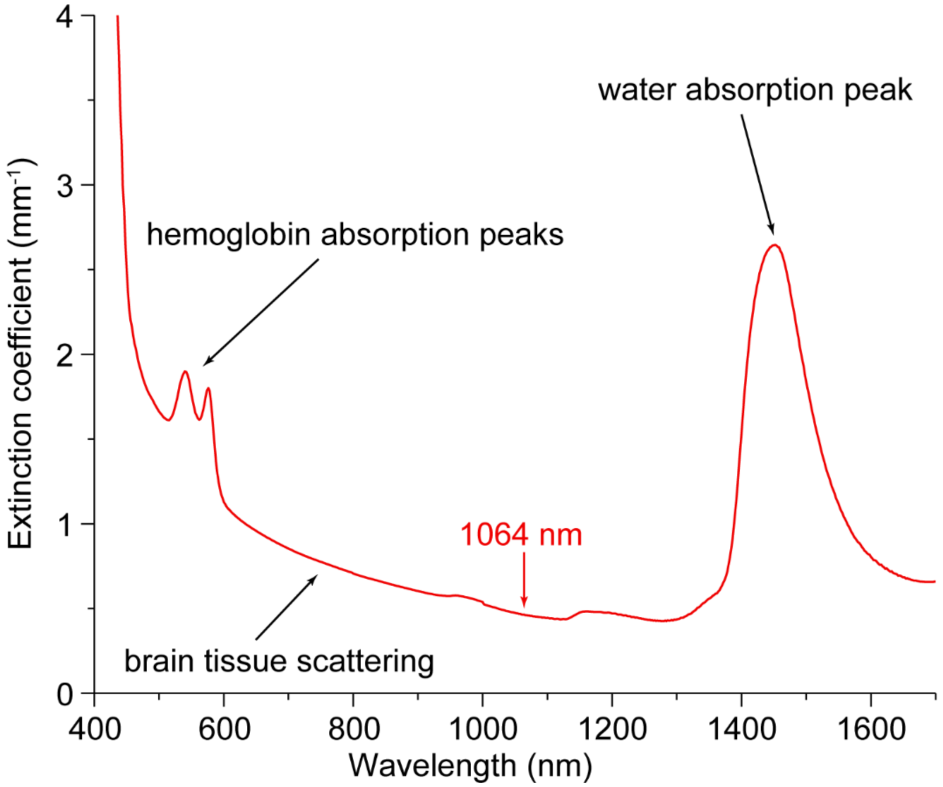


**Supplementary Fig. 1**. Simulated extinction coefficient spectrum of brain tissue from 400 to 1700 nm. This spectrum is calculated based on the reported brain scattering coefficient[11], water absorption coefficient, oxyhemoglobin and deoxyhemoglobin absorption coefficients weighted by the composition of water and hemoglobin in the brain tissue[12]. The main contributions from the water absorption, hemoglobin absorption and brain tissue scattering are labelled. The accuracy of this extinction spectrum is validated by comparing it to experimental measurements at selected wavelengths[22].

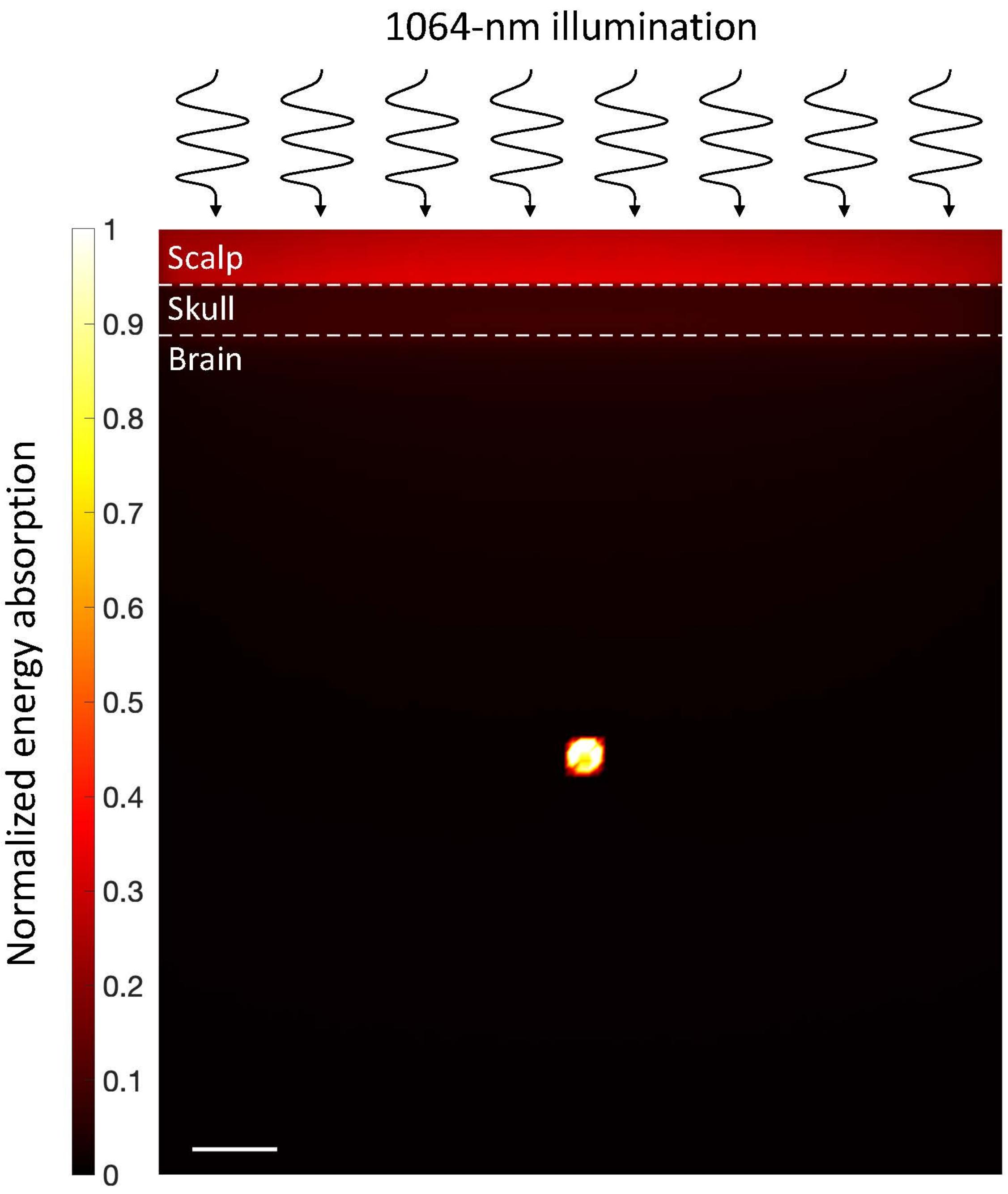


**Supplementary Fig. 2.** Monte Carlo simulation of energy absorption by the scalp, skull, brain tissue, and MINDS injected in the brain upon 1064-nm illumination above the scalp. 1.8 mg $mL^{-1}$ of MINDS is confined in a 400 μm diameter circle located 5 mm below the surface of the brain (i.e., 6.3 mm below the surface of the scalp). MINDS absorb more photon energy than any other tissues (indicated as the "hot spot" in the brain) despite being located deep in the brain, owing to the deep tissue penetration of 1064-nm light and the strong absorption of 1064-nm light by MINDS. The scale bar indicates 1 mm.

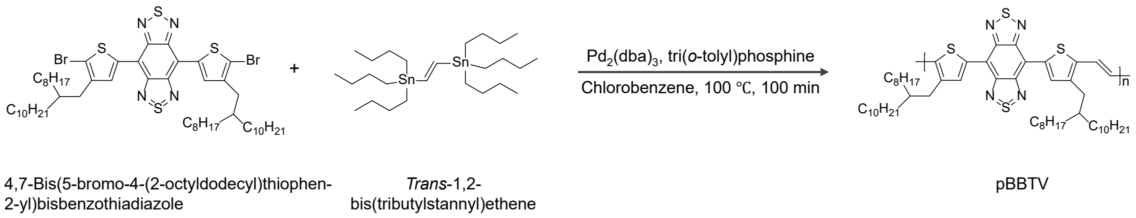


**Supplementary Fig. 3.** Synthetic route of pBBTV.

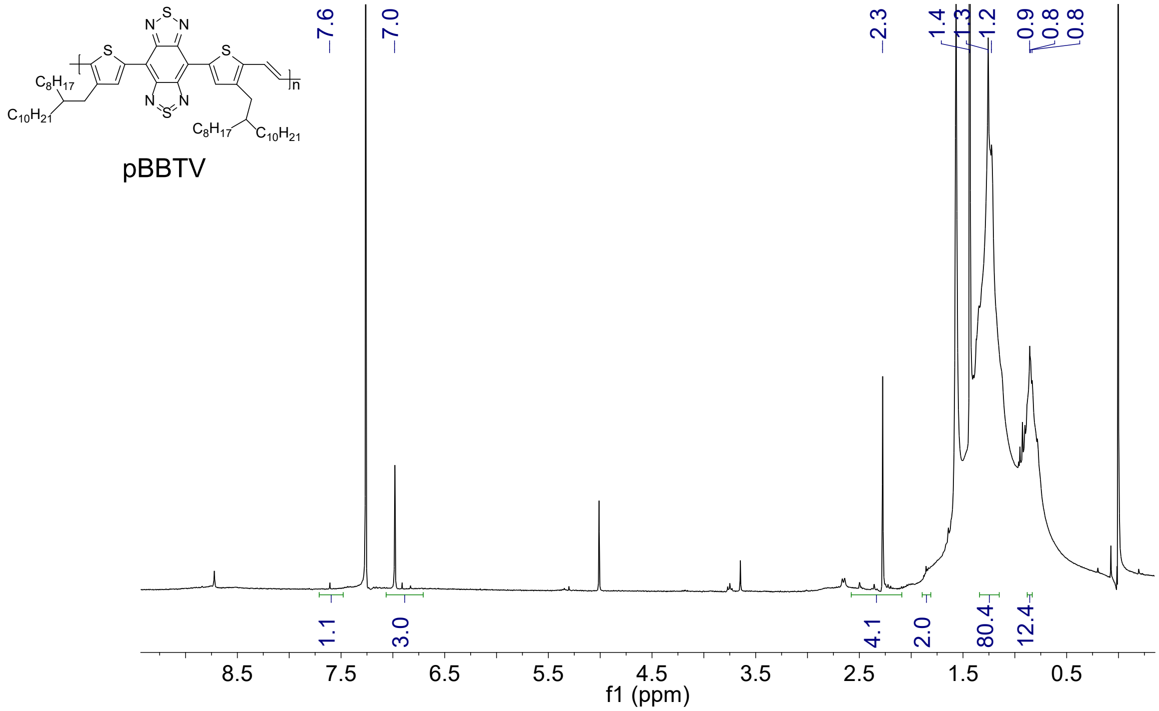


**Supplementary Fig. 4.** $^{1}$H NMR spectrum of pBBTV in $CDCl_3$.

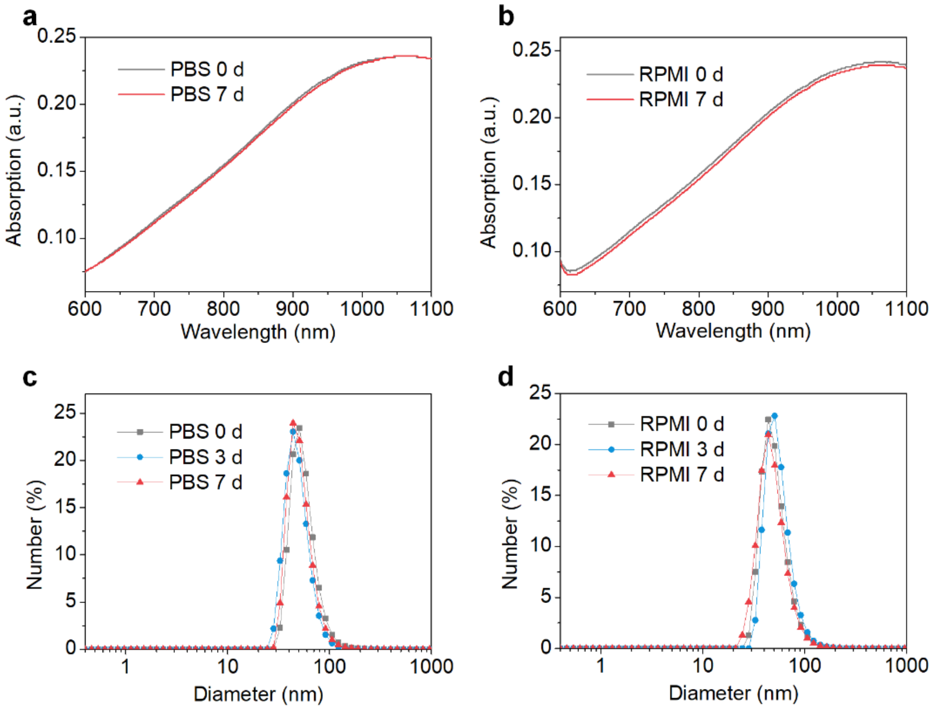


**Supplementary Fig. 5.** Structural stability of MINDS under physiological conditions. **(a & b)** Absorption spectra of MINDS after incubation in 1× PBS **(a)** and RPMI 1640 complete culture medium (RPMI) **(b)** at 37 ºC for 7 days. **(c & d)** Dynamic light scattering profiles of MINDS after incubation in 1× PBS **(c)** and RPMI **(d)** at 37 ºC for 3 or 7 days.

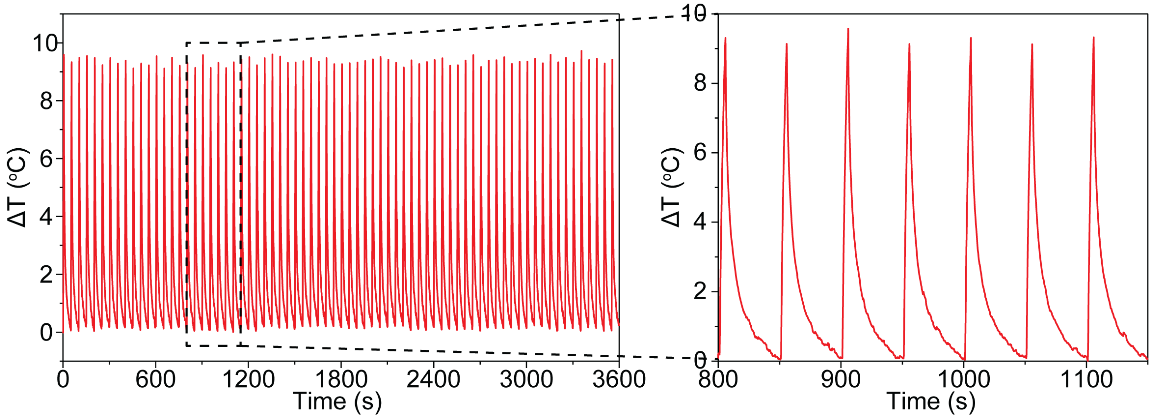


**Supplementary Fig. 6.** *In vivo* photostability of MINDS demonstrates repeated photothermal heating and cooling for at least 1 h. A 1064 nm laser with a power density of 8 mW mm$^{-2}$ was used to illuminate the mouse cortex injected with a MINDS solution of 1.8 mg mL$^{-1}$ and a thermal camera was used to monitor the temperature of the cortex.

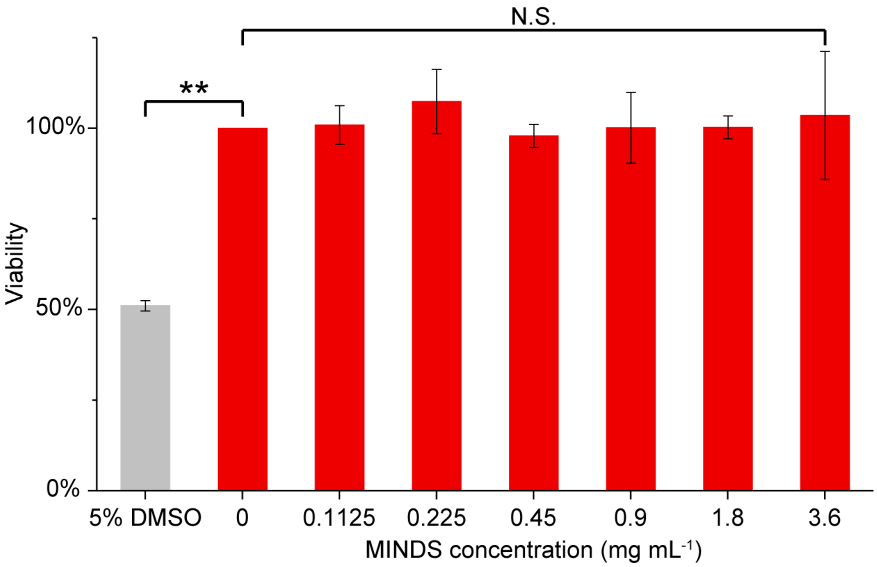


**Supplementary Fig. 7.** Cytotoxicity assessment of MINDS. Cell viability assay reveals minimum toxicity of MINDS up to 3.6 mg $mL^{-1}$ in primary neuron culture (N.S., not significant). As a comparison, MINDS concentration used for *in vivo* stereotaxic injection was 1.8 mg $mL^{-1}$. In contrast, the viability of the positive control group, which is comprised of cells incubated with 5% DMSO dropped to ca. 50% (**, $P < 0.01$). Error bars indicate SD ($n = 3$ per concentration group).

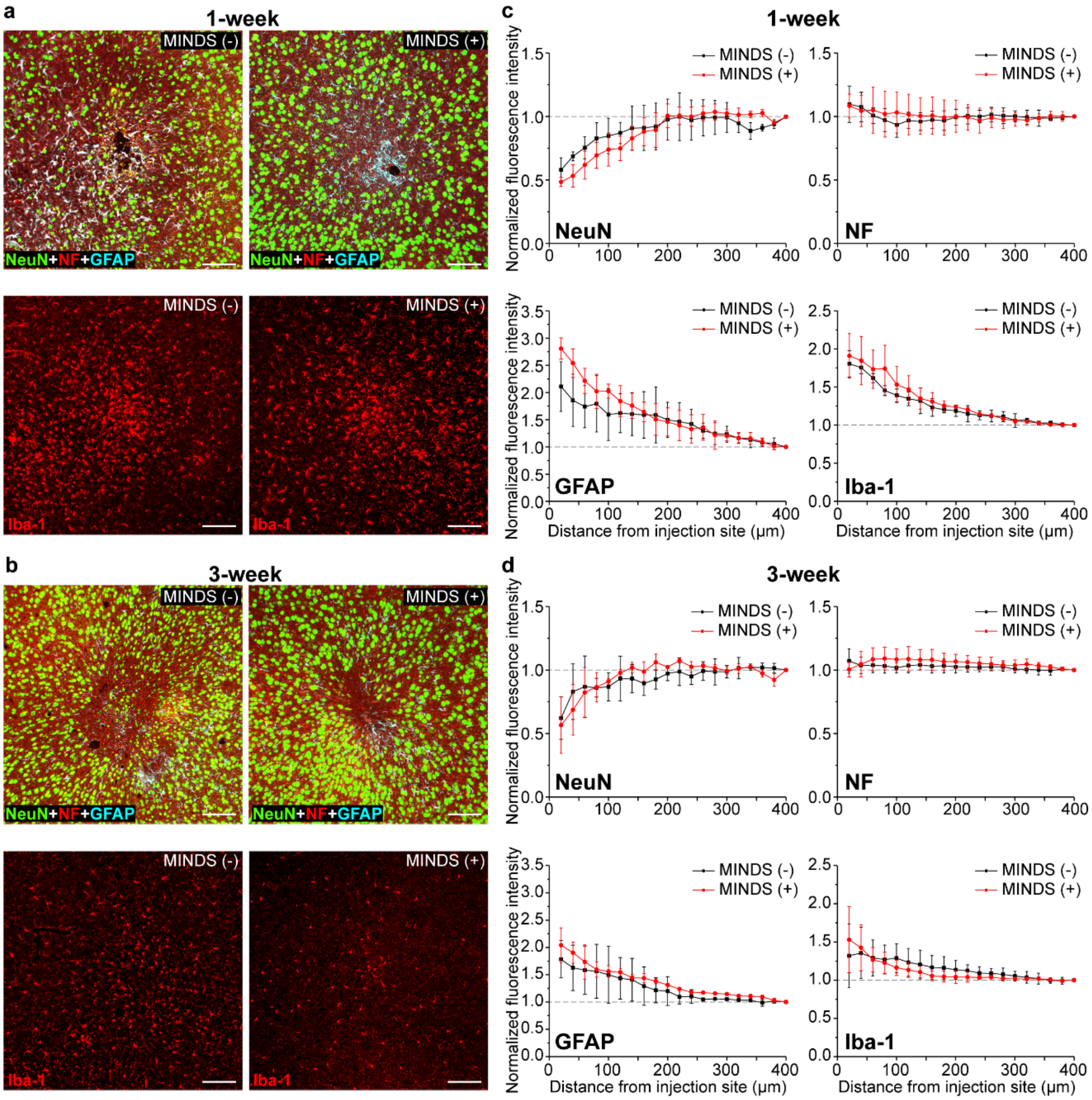


**Supplementary Fig. 8.** Assessment of the chronic immune response in the mouse brain due to MINDS injection. (**a&b**) Confocal fluorescence images characterizing the immune response near the injection sites of MINDS or the carrier at 1 week **(a)** or 3 weeks **(b)** post injection. In the top panels, neuron nucleus (NeuN), neurofilaments (NF; indicating neurites) and glial fibrillary acidic protein (GFAP; indicating astrocytes) are labeled in green, red and cyan, respectively. In the bottom panels, ionized calcium binding adaptor molecule 1 (Iba1; indicating microglia) is labelled in red. The scale bars indicate 100 μm. (**c&d**) Normalized fluorescence intensity of NeuN, NF, GFAP and Iba-1 as a function of distance from the MINDS/carrier injection sites at 1 week **(c)** and 3 weeks **(d)** post injection. Error bars indicate SD (*n* = 3 per group).

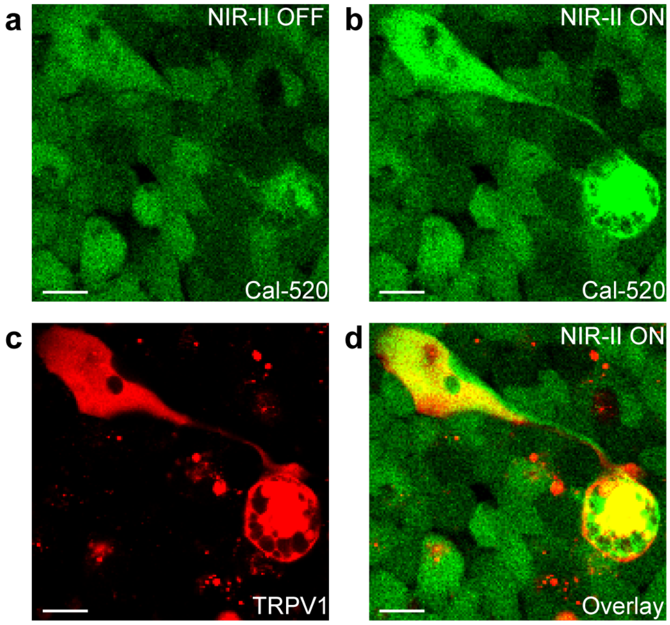


**Supplementary Fig. 9.** TRPV1 expression is colocalized with the calcium signal increase under NIR-II photothermal stimulation. (**a** & **b**) Confocal fluorescence images of HEK293T cells transfected with pAAV-CMV-TRPV1-mCherry before (**a**) and after NIR-II photothermal stimulation (**b**) in the Cal-520 channel. (**c**) Confocal fluorescence image of the same cells in the mCherry channel, showing the expression of TRPV1. (**d**) An overlay of **b** and **c**, showing the colocalization of calcium signal increase and TRPV1 expression in the same cells. The scale bars indicate 20 μm.

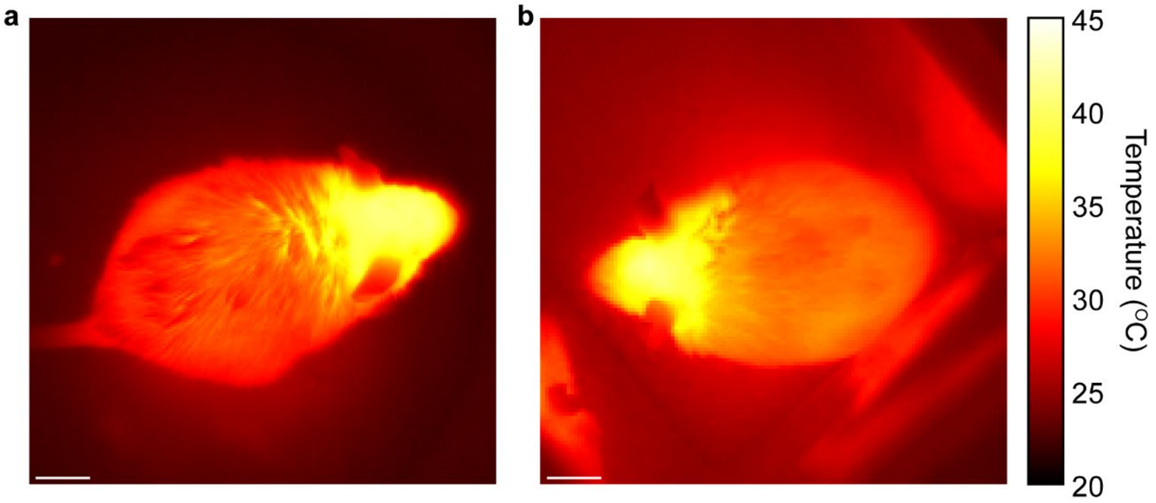


**Supplementary Fig. 10.** Thermal images of mice under 1064-nm illumination during the unilateral circling test (**a**) and the conditioned place preference test (**b**). The scale bars indicate 1 cm.

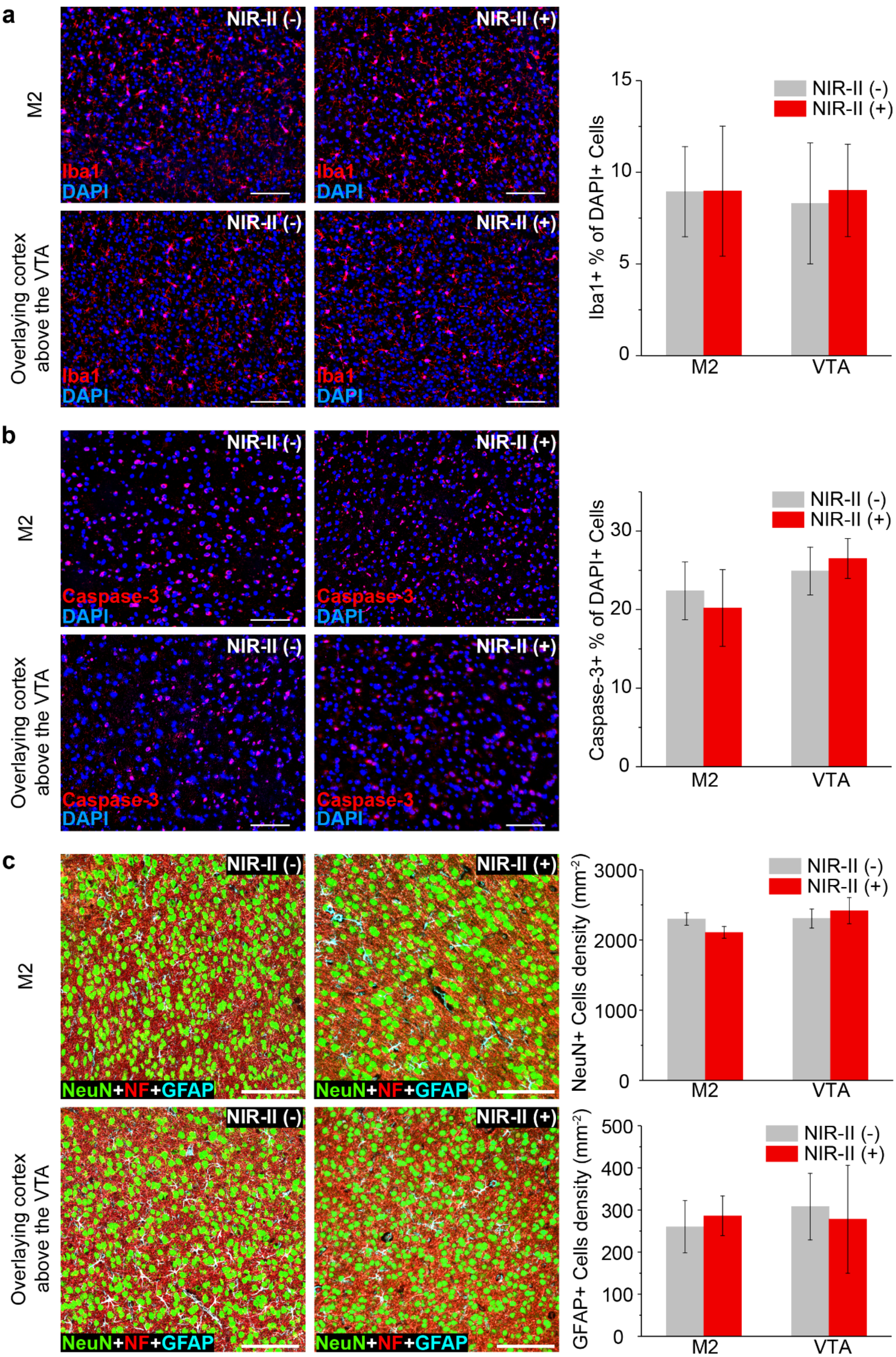
a
M2
Overlaying cortex above the VTA
NIR-II (-)
NIR-II (+)
Iba1
DAPI
Iba1+ % of DAPI+ Cells
M2
VTA
b
Caspase-3
DAPI
Caspase-3+ % of DAPI+ Cells
c
NeuN+NF+GFAP
NeuN+ Cells density (mm⁻²)
GFAP+ Cells density (mm⁻²)

**Supplementary Fig. 11.** Assessment of thermal damage through the immunohistochemical labeling of Iba1 (**a**), cleaved caspase-3 (Caspase-3; indicating apoptosis) (**b**), NeuN, NF and GFAP (**c**). In each panel, mice were injected with MINDS in M2 or VTA, treated with or without the NIR-II illumination protocol used in the corresponding behavioral test, and sacrificed 1 week after the treatment. The left panels are confocal images of different biomarkers in M2 (top) or the cortex region above VTA (bottom) with (right) or without (left) NIR-II illumination. The cortex above VTA was chosen as the worst-case scenario to evaluate the nonspecific thermal damage to the neural tissue due to its closest distance to NIR-II illumination. The scale bars indicate 100 μm. In **a&b**, the right panels are the percentages of Iba1+ and Caspase-3+ cells within cell population indicated by DAPI, respectively. In **c**, the right panels are NeuN+ (top) or GFAP+ (bottom) cell densities. All error bars indicate SD ($n$ = 3 for all experiments). No statistically significant difference is found for any pairwise comparisons in the bar charts of the right panels.

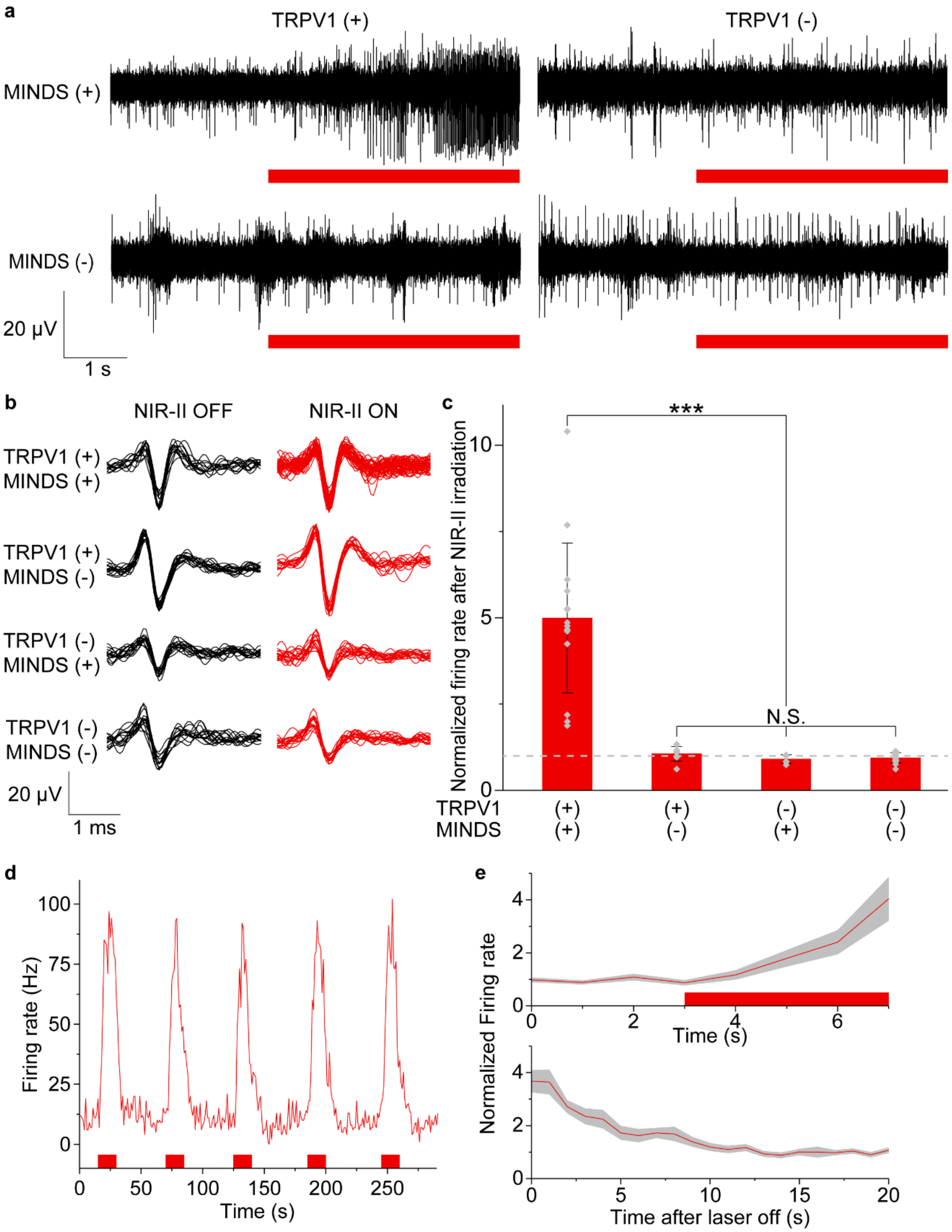


**Supplementary Fig. 12.** *In vivo* electrophysiological recording in the mouse hippocampus with NIR-II neuromodulation. **(a)** Representative traces for the experiment group and the control groups with NIR-II illumination. **(b)** Representative overlaid spike waveforms during the same duration before and after NIR-II illumination for each group. **(c)** Statistical analysis of the normalized change in the neuron firing rate upon NIR-II illumination for animals under different experiment conditions. Only the TRPV1+/MINDS+ animals showed a statistically significant increase in neuron firing rate compared to the control groups (***, $P < 0.001$). The gray dashed line indicates a normalized firing rate of 1. The firing rates are normalized to account for the intrinsically different spontaneous neuron activities in different measurements. The error bars indicate SD, and each point indicates an independent trial. (**d**) Representative neural firing rate dynamics showing reproducible control of neural activities by NIR-II illumination. (**e**) The average onset (top) and offset (bottom) dynamics of normalized neural firing rate upon and after NIR-II stimulation, respectively. The shade indicates ±1 standard error of the mean from $n = 23$ traces. The red bars in **a, d & e** indicate the duration of NIR-II illumination. For the bottom panel in **e**, NIR-II illumination was turned off at $t = 0$ s.

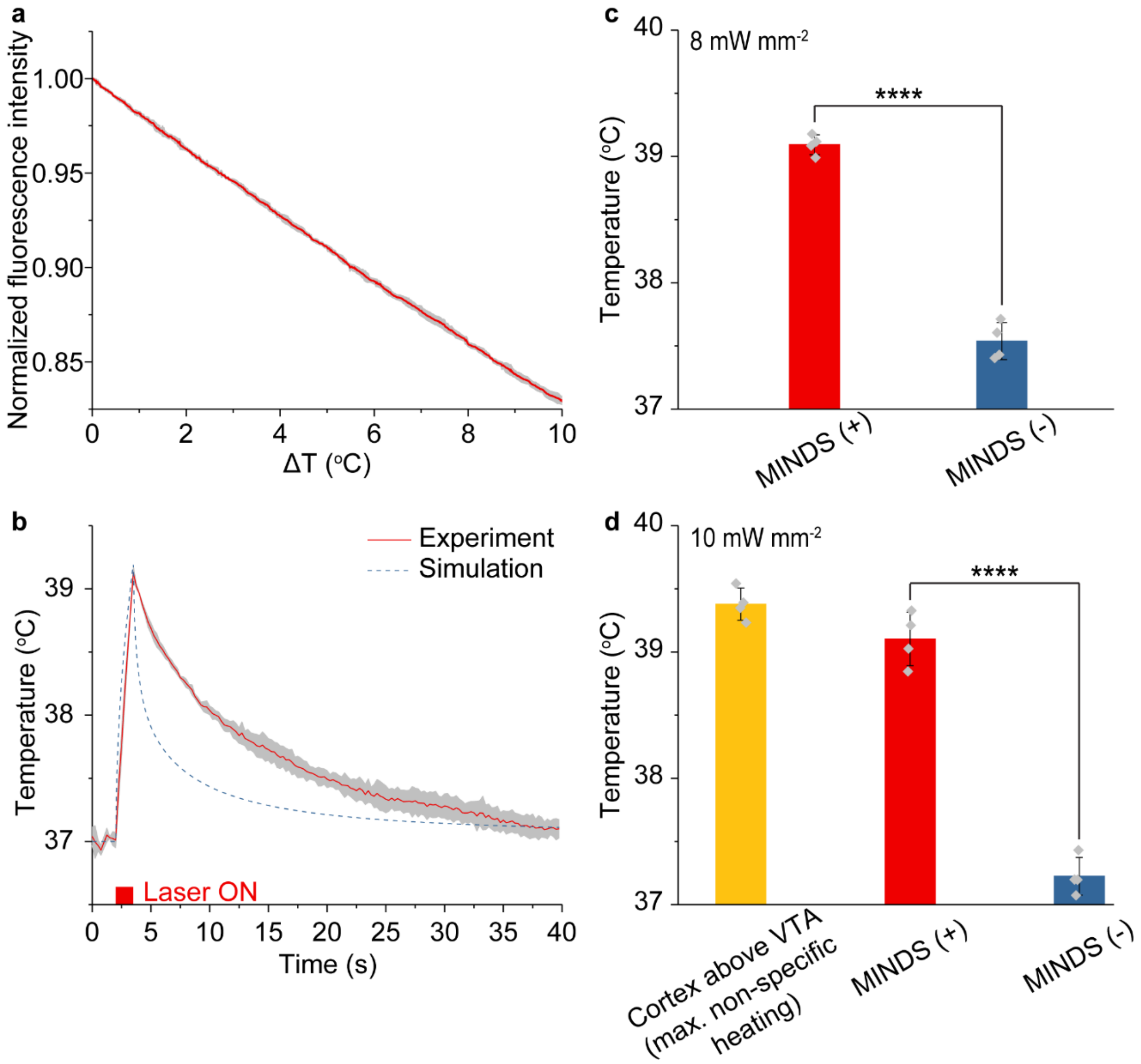


**Supplementary Fig. 13.** *In vivo* temperature measurements based on temperature dependent fluorescence intensity of DyLight550. (**a**) Calibration of the temperature dependence of the fluorescence intensity of DyLight550. The percentile change of fluorescence intensity per degree of temperature increase was calculated to be -1.71% $^{o}C^{-1}$. (**b**) Temperature dynamics of M2 in the presence of MINDS under 8 mW $mm^{-2}$ NIR-II illumination. The red curve is the experimentally measured data while the blue dashed curve is the prediction from Monte Carlo stimulation with thermal diffusion. (**c**) The final temperatures in M2 with or without MINDS after 1.5 s of 8 mW $mm^{-2}$ NIR-II illumination. The MINDS (+) group showed a significantly higher temperature increase than the MINDS (-) group (****, $P < 0.0001$). (**d**) The final temperatures in the VTA with or without MINDS, along with the temperature in the cerebrum cortex above the VTA after 5.5 s of 10 mW $mm^{-2}$ NIR-II illumination. The temperature in the cortex above the VTA sets the upper limit of non-specific heating in the brain tissue since NIR-II light illuminates from above. The MINDS (+) group showed a significantly higher temperature increase than the MINDS (-) group (****, $P < 0.0001$). The shades (in **a&b**) and the error bars (in **c&d**) indicate ±1 SD from $n = 4$ trials.

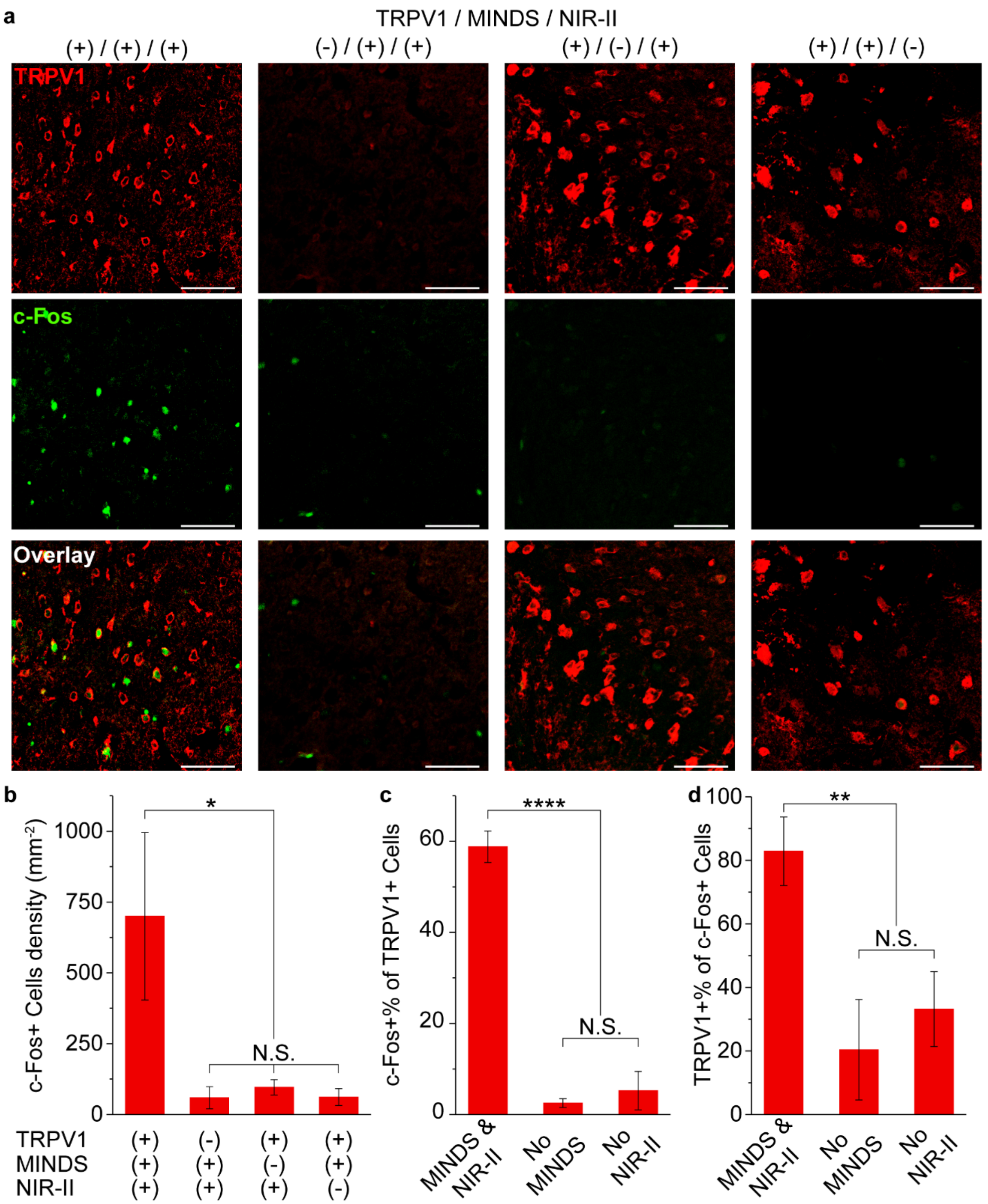


**Supplementary Fig. 14.** Immunohistology of the M2 region after NIR-II illumination. (**a**) Confocal images of the M2 region under different experiment conditions. An increase in c-Fos expression driven by NIR-II stimulation was only observed in the presence of both TRPV1 and MINDS. The scale bars indicate 50 μm. (**b**) Statistical analysis for the density of c-Fos+ cells for the four experiment conditions in **a**. (**c**) Percentage of c-Fos+ cells within the TRPV1+ cell population. (**d**) Percentage of TRPV1+ cells within the c-Fos+ cell population. Both **c&d** were analyzed for the three experimental groups with TRPV1 transduction. In **b-d**, a statistically significant difference is found for the triple-positive group (TRPV1+, MINDS+, and NIR-II+) vs. all other conditions (*, $P < 0.05$; **, $P < 0.01$; ****, $P < 0.0001$). All other groups do not have any statistically significant differences in pairwise comparisons (N.S., not significant). All error bars indicate SD ($n = 3$ per group).

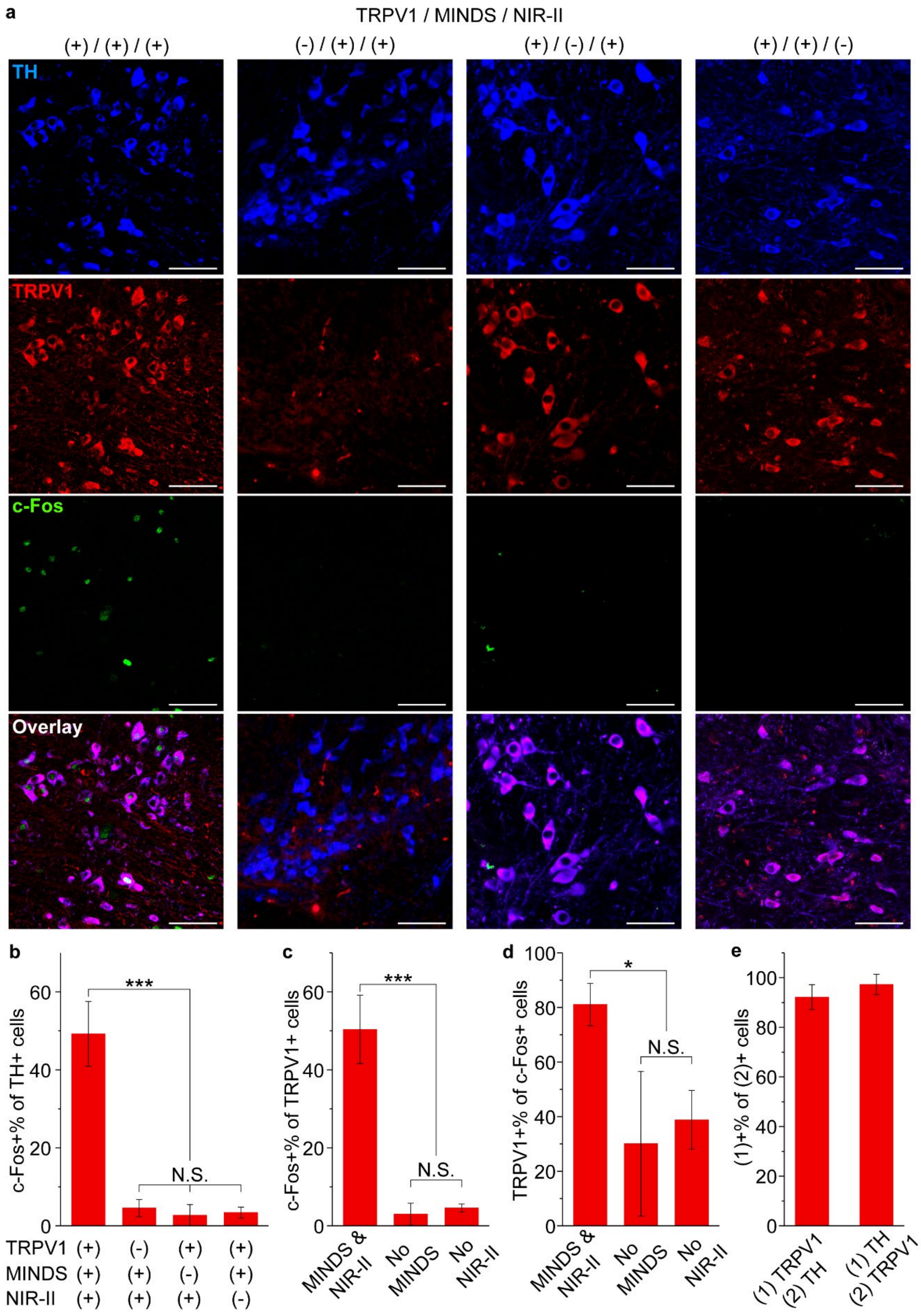
a
TRPV1 / MINDS / NIR-II
(+) / (+) / (+)
(-) / (+) / (+)
(+) / (-) / (+)
(+) / (+) / (-)
TH
TRPV1
c-Fos
Overlay
b
c-Fos+% of TH+ cells
***
N.S.
TRPV1 (+) (-) (+) (+)
MINDS (+) (+) (-) (+)
NIR-II (+) (+) (+) (-)
c
c-Fos+% of TRPV1+ cells
***
N.S.
MINDS & NIR-II
No MINDS
No NIR-II
d
TRPV1+% of c-Fos+ cells
*
N.S.
MINDS & NIR-II
No MINDS
No NIR-II
e
(1)+% of (2)+ cells
(1) TRPV1 (2) TH
(1) TH (2) TRPV1

**Supplementary Fig. 15.** Immunohistology of the VTA region after NIR-II illumination. (**a**) Confocal images of the VTA region under different experimental conditions. An increase in c-Fos expression driven by NIR-II stimulation was only observed in the presence of both TRPV1 and MINDS. The scale bars indicate 50 μm. (**b**) Percentage of c-Fos+ neurons in the TH+ cell population, corresponding to the four conditions presented in **a**. (**c**) Percentage of c-Fos+ cells within the TRPV1+ cell population. (**d**) Percentage of TRPV1+ cells within the c-Fos+ cell population. Both **c&d** were analyzed for the three experimental groups with TRPV1 transduction. In **b-d**, a statistically significant difference is found for the triple-positive group (TRPV1+, MINDS+, and NIR-II+) vs. all other conditions (*, $P < 0.05$; ***, $P < 0.001$). All other groups do not have any statistically significant differences in pairwise comparisons (N.S., not significant). (**e**) Statistics of TRPV1 expression in TH+ neurons (left bar) and TH expression in TRPV1+ neurons (right bar). All error bars indicate SD ($n = 3$ per group).

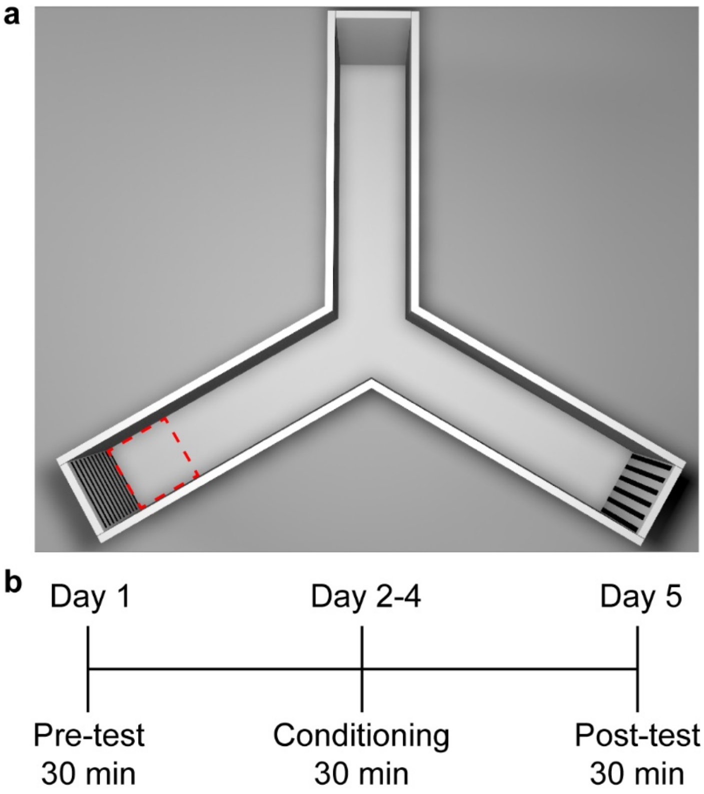


**Supplementary Fig. 16.** Setup and timeline for conditioned place preference test. (**a**) 3D scheme showing the context at the end of each arm of the Y-maze. The red dashed square indicates the region where NIR-II light is illuminated. (**b**) Timeline of mouse place preference test in the Y-maze.

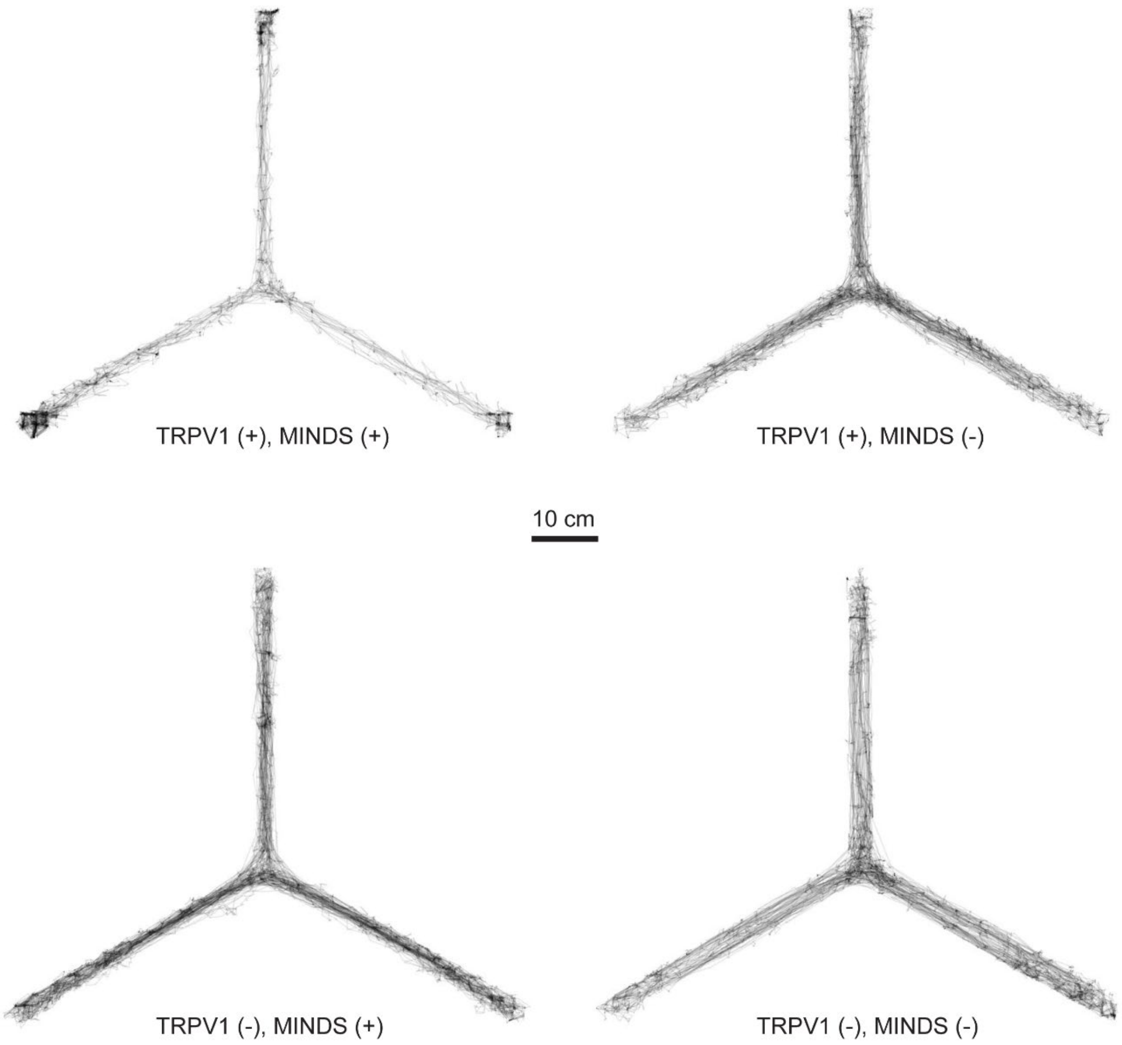


**Supplementary Fig. 17.** Post-test mice trajectories under different experiment conditions, corresponding to the heat maps in **Fig. 4d**.

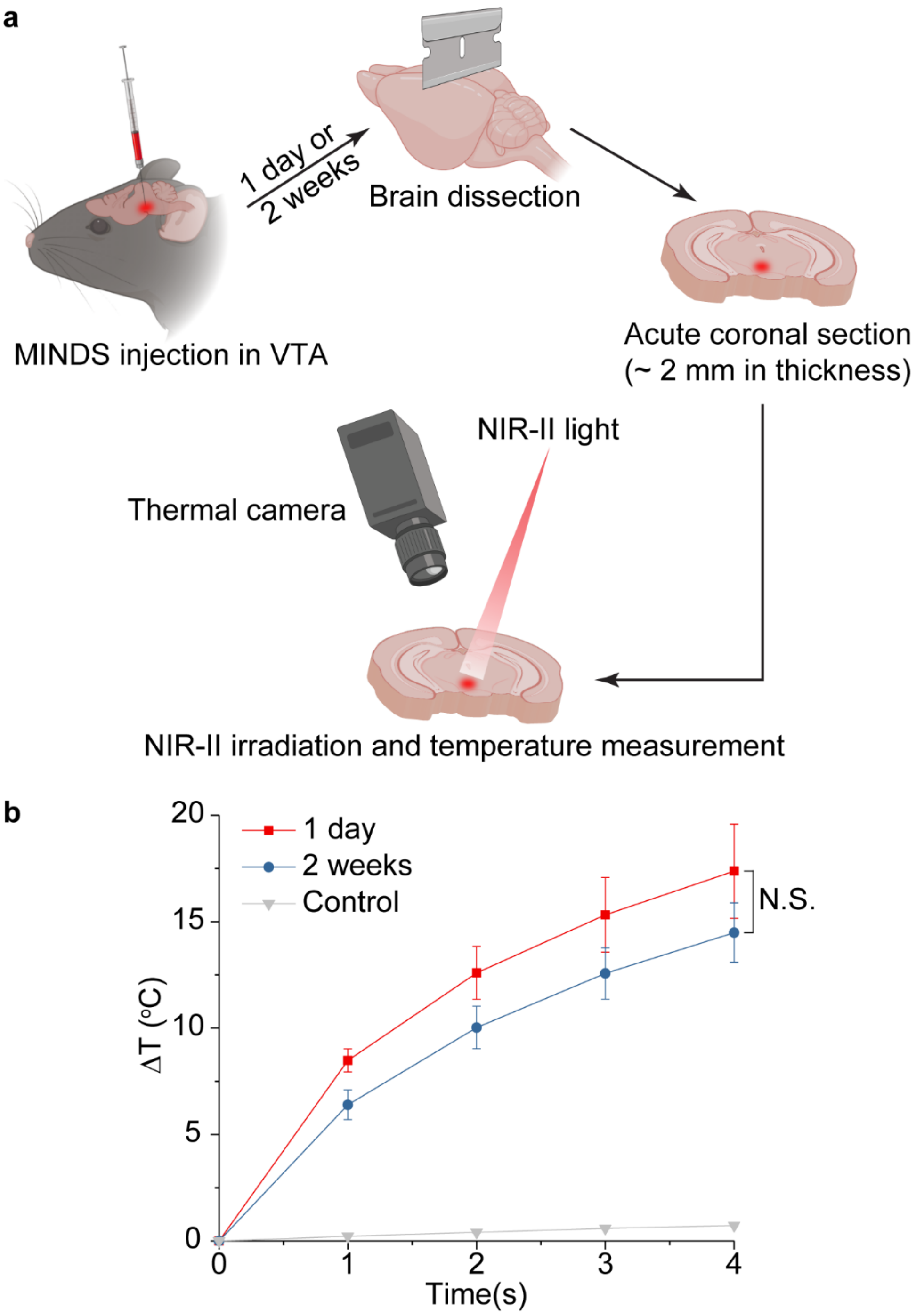


**Supplementary Fig. 18.** *In vivo* chronic functional stability of MINDS. **(a)** Schematics showing the experiment setup for assessing the chronic utility of MINDS *in vivo*. (**b**) Temperature dynamics of VTA slices at 1 day or 2 weeks post injection upon 10 mW $mm^{-2}$ NIR-II light. After 4 s of heating, the temperature increase for the 2-week group does not have a statistically significant difference from that of the 1-day group (N.S., not significant). The temperature dynamics of the control group were measured by placing MINDS-free brain slices under NIR-II illumination with the same power density and duration. The error bars indicate SD ($n$ = 3 for all groups).

**Supplementary Table 1.** Summary of the deleterious consequences of tethering.

| Deleterious consequences of tethering | References |
|---|---|
| "*However,* ***tethered experiment setups present several problems for the experimenter****. Animals must be handled at the beginning of each experiment, which can* ***alter behavior****.* ***Tethering may be impractical for optogenetic experiments over long periods of time*** *such as desired in many developmental, longitudinal, disease-progression, or other chronic-perturbation experiments, where* ***fibers and cables risk breakage or binding over time****. Finally****, the need for tethering sets limits on the number of animals that can be manipulated in a single experiment*** *(e.g.,* ***a social experiment with multiple mice could result in tangling or breakage of the tethers****) or in parallel experiments (e.g., in high throughput screening of large sets of animals).*" | *J. Neural Eng.* **8**, 046021 (2011) |
| "***Tethering of the animal is one particular limitation of current optogenetic systems.*** *Tethering increases experimental risks due to tangling of the optical fiber cord around the animal or through the experience of unintended rotational forces when connected.* ***These issues confound natural behaviors that are essential for interpreting behavioral data.***"<br>"*In the optogenetic technique,* ***tethering the animal*** *to the benchtop light source and control system substantially* ***reduces the reproducibility of validated behavioral tests****.*"<br>"*Despite achieving rotary optimization,* ***tethering can still alter behavior and disrupt the naturalistic testing environment*** *being sought, potentially* ***confounding behavioral experiments****.*" | *Neurophotonics* **2**, 031206 (2015) |
| "*Despite the success of these studies,* ***tethered fiber-optic approaches*** *have* ***restricted opportunities for the study of more complex, ethologically relevant behavioral paradigms*** *such as enclosed home-cage behavior, spontaneous pain, wheel running and freely moving social interactions.*" | *Nat. Protoc.* **8**, 2413–2428 (2013) |
| "*These* ***tethered systems*** *nonetheless* ***impose significant constraints on experimental design and interpretation****, both by* ***requiring investigators to*** | *Nat. Methods* **12**, 969–974 (2015) |

| | |
|---|---|
| ***handle and physically restrain animals to attach an optical fiber*** *before behavioral testing and by* ***limiting the environments in which optogenetic experiments can be performed****.*" "*However, these* ***tethered devices are typically bulky and restrict animal movement****.*" | |
| "*The main advantage imparted by biotelemetry over conventional methods is the elimination of* ***confounding stress effects*** *introduced by handling,* ***restraint****, and anesthesia.*" | *Am. J. Physiol. Regul. Integr. Comp. Physiol.* **286**, R967-R974 (2004) |
| "*Moreover, in certain conditions the* ***need for a tether*** *or implanted telemetry device* ***can potentially affect the behavior of the animal****, particularly in mice where the implant size is typically disproportionately large compared to body mass.*" | *J. Bio. Rhythms* **27**, 48-58 (2012) |
| "*Another major breakthrough can be expected through the development of* ***wireless communications and wireless powering of the electrophysiological and optogenetic recording and delivery devices in freely moving animals****. There are several excellent bioengineering labs that are pushing the envelope on this key technology that will* ***make the need for tethered in vivo recordings from behaving animals history****. The need is clear; for example, in my lab, where we do 24/7 video-EEG monitoring and closed-loop optogenetics in mice,* ***the tethered nature of the recordings is a huge challenge, especially since the animals are having robust behavioral seizures****, so it would be truly transformative to have wireless recording and light-powering technology that would negate the need for* ***long wires and optical cables that can twist and break****.*" | *Nat. Neurosci.* **18**, 1202–1212 (2015) |
| "*Thus,* ***unexpected caloric intake differences*** *between the previous and current studies following the DS feeding paradigm may be due to housing conditions (****tethered EEG recording cables*** *vs homecage locomotor boxes).*" "*... mice* ***did not experience normal weight gain****. We believe this was* ***due to the combined stressors*** *required for each signal acquisition (****tether cables for EEG****...*" | *PLoS One* **13**, e0196743 (2018) |

**Supplementary Table 2.** Summary of reported photothermal conversion efficiencies at 1064 nm.

| Material | Photothermal conversion efficiency η (%) | Reference |
|---|---|---|
| AuPB | 80.8 | *ACS Nano* **12,** 2643-2651 (2018). |
| PEDOT:ICG@PEG-GTA | 71.1 | *Biomaterials* **41,** 132-140 (2015) |
| MINDS | 71 | This study |
| RBC@$Cu_{2-x}$SeNPs | 67.2 | *Chem. Commun.* **55,** 6523-6526 (2019). |
| $Fe_3O_4$@PPy@GOD NCs | 66.4 | *Adv. Mater.* **31,** 1805919 (2019). |
| $NP_{PBTPBF-BT}$ | 66.4 | *Biomaterials* **155,** 103-111 (2018). |
| Gold Nanoraspberry | 65 | *Chem. Commun.* **55,** 4055-4058 (2019) |
| Ultrathin polypyrrole nanosheets | 64.6 | *Nano Lett.* **18,** 2217−2225 (2018) |
| NiP PHNPs | 56.8 | *Nano Lett.* **19**, 5093-5101(2019) |
| $Fe_2P$ | 56.6 | *Angew. Chem. Int. Ed.* **58,** 2407 –2412 (2019) |
| SPN-PT | 53 | *Adv. Mater.* **31,** 1808166 (2019). |
| TBDOPV-DT | 50.5 | *Chem. Mater.* **29**, 718-725 (2017) |
| $Ti_2O_3$@HA NPs | 50.2 | *J. Mater. Chem. B* **6,** 7889-7897 (2018) |
| TBDOPV-DT NPs | 50 | *ACS Appl. Mater. Interfaces* **10,** 7919−7926 (2018). |
| DPP-IID-FA NPs | 49.5 | *Chem. Commun.* **54,** 13599-13602 (2018). |
| SPN-DT | 49 | *Adv. Mater.* **31,** 1808166 (2019). |
| H-$SiO_x$ NPs | 48.6 | *Biomaterials* **143,** 120-129 (2017). |

| | | |
|---|---|---|
| P3 NPs | 46 | *ACS Nano* **13,** 7345−7354 (2019) |
| $Nb_2C$ NS | 45.7 | *J. Am. Chem. Soc.* **139,** 16235−16247 (2017) |
| $V_2C$-TAT@Ex-RGD | 45.1 | *ACS Nano* **13,** 1499-1510 (2019). |
| PEG-TONW NRs | 43.6 | *Nano Lett.* **19,** 1179-1189 (2019). |
| $SPN_{I-II}$ | 43.4 | *Adv. Mater.* **30,** 1705980 (2018). |
| Bi@C NPs | 43.2 | *Nanoscale* **11,** 9906-9911 (2019) |
| Au NPL@$TiO_2$ | 42.1 | *Nanoscale* **11,** 2374–2384 (2019). |
| $Cu_3BiS_3$ NR | 40.7 | *Biomaterials* **112,** 164-175 (2017) |
| $TiO_2$@$TiO_{2-x}$ | 39.8 | *ACS Nano* **12,** 4545–4555 (2018). |
| $Ti_3C_2$@Au | 39.6 | *ACS Nano* **13,** 284−294 (2019) |
| $(NH_4)_xWO_3$ | 39.2 | *Biomaterials* **52,** 407-416 (2015). |
| Platinum nanoworms | 38.9 | *J. Mater. Chem. B* **6,** 5069-5079 (2018) |
| PEG-$MoO_x$ | 37.4 | Nanoscale **10,** 1517–1531 (2018). |
| Au-$Cu_9S_5$NPs | 37 | *J. Am. Chem. Soc.* **136,** 15684-15689 (2014). |
| SPN-OT | 36 | *Adv. Mater.* **31,** 1808166 (2019). |
| Bi-LyP-1 | 32.2 | *ACS Nano* **11,** 3990-4001 (2017). |
| PtNPs | 30.9 | *Biomaterials* **34,** 5833-5842 (2013). |
| PVP-$Cu_3BiSe_3$ NPs | 30.7 | *Nanoscale* **11,** 7157–7165 (2019). |
| IONP@shell-in-shell NPs | 28.3 | *ACS Appl. Mater. Interfaces* **10,** 1508−1519 (2018). |
| PEG−CuS−Au−MnO2 ternary JNPs | 28.0 | *ACS Appl. Mater. Interfaces* **10,** 24137−24148 (2018). |
| $Fe_3O_4$@CuS-PEG | 19.2 | *Adv. Funct. Mater.* **25,** 6527–6537 (2015). |

**Supplementary Table 3.** Parameter comparison for emerging neural stimulation methods.

| Stimulation modalities | System(s) of study | Invasiveness for *in vivo* study | *In vivo* cell-type specificity (+/-)* | Medium for wireless stimulation (size) | Power density (wavelength) | Temporal response | Reference |
|---|---|---|---|---|---|---|---|
| Photothermal | Mouse behavioral study and cells | Through scalp and skull | + | Nanoparticles (~40 nm) | 8-10 mW $mm^{-2}$ (1064 nm, mouse) and 400 mW $mm^{-2}$ (1040 nm, cell) | 5.0±1.5 s (mouse) and 1.1±0.2 s (cell) | This study |
| | Cells | N/A | N/A | Nanoparticles (25-37 nm) | ~$1.0\times10^5$ mW $mm^{-2}$ (808 nm) | ~0.1 s | *J. Am. Chem. Soc.* **138**, 9049-9052 (2016). |
| | Cells | N/A | N/A | Nanorods (size not reported) | $8\times10^3$ mW $mm^{-2}$ (780 nm) | N/A | *Angew. Chem. Int. Ed.* **54**, 11725-11729 (2015). |
| | Cells and brain slice | N/A | N/A | Nanoparticles (~20 nm) | ~$3.1\times10^5$ mW $mm^{-2}$ (532 nm) | ~ms | *Neuron* **86**, 207-217 (2015). |
| | Cells | N/A | N/A | Mesostructured silicon (1-2 μm) | ~$6.8\times10^4$ mW $mm^{-2}$ (532 nm) | ~ms | *Nat. Mater.* **15**, 1023-1030 (2016). |
| | Cells | N/A | N/A | Water molecules | 0.73-3.8 W (1869 nm & 1889 nm) | ~ms | *Nat. Commun.* **3**, 736 (2012). |
| | Cells | N/A | N/A | Water molecules | ~$1.0\times10^6$ mW $mm^{-2}$ (980 nm) ~$3.8\times10^5$ mW $mm^{-2}$ (1460 nm) | ~ms | *Biophys. J.* **96,** 3611-3619 (2009) |

| | | | | | | | |
|---|---|---|---|---|---|---|---|
| | Anesthetized mouse, hydra and brain slice | Scalp and skull removed | - | Nanoparticles (30-40 nm) | 4-40 mW (1040 nm) | ~1 s (mouse) and 0.1-0.5 s (cell) | *Light Sci. Appl.* **7**, 100 (2018). |
| Optical and optogenetics | Mouse behavioral study | Subretinal injection | + | Nanoparticles (38 nm±2 nm) | $1.62 \times 10^{-2}$ mW $mm^{-2}$ (980 nm) | ~ms | *Cell* **177,** 243-255 (2019) |
| | Mouse behavioral study | Chronic brain implant | + | Optoelectronic device (~ 15 µm × 400 µm × 1 mm) | 5-23.5 mW $mm^{-2}$ (450 nm) | ~ms | *Science* **340**, 211-216 (2013). |
| | Head-fixed mice | Scalp removed with intact skull | + | Red-shifted opsins, Jaws | 127 mW $mm^{-2}$ (635 nm, mouse) | ~ms | *Nat. Neurosci.* **17,** 1123-1129 (2014). |
| | Head-tethered mice | Scalp removed with intact skull | + | Red-shifted opsin, ChRmine | Up to 3200 mW $mm^{-2}$ (635 nm, mouse) | ~ms | *Nat. Biotechnol.* DOI: 10.1038/s41587-020-0679-9 (2020) |
| | Head-tethered mice and head-fixed monkeys | Scalp removed with intact skull (mouse); Scalp and skull removed with intact dura (monkey) | + | Ultrasensitive, step-function opsin, SOUL | 50 mW (473 nm, mouse)<br><br>Up to 173 mW $mm^{-2}$ (473 nm, monkey) | ~s | *Neuron* **107,** 38-51 (2020) |
| | Head-tethered mice | Scalp removed with intact skull | + | Ultrasensitive opsin, ChRger2 | 20 mW (447 nm, mouse) | ~ms | *Nat. Methods* **16,** 1176-1184 (2019) |
| | Mouse behavioral study and cells | Scalp removed with intact skull | + | Nanoparticles (~90 nm) | 1400 mW $mm^{-2}$ (980 nm, mouse) and 350-8220 mW $mm^{-2}$ (980 nm, cell) | ~ms | *Science* **359**, 679-684 (2018). |
| | Mouse behavioral study | Scalp removed with intact skull | + | Optoelectronic device (10-25 $mm^3$) | 10-60 mW $mm^{-2}$ (470 nm) | ~ms | *Nat. Methods* **12**, |

| | | | | | | | |
|---|---|---|---|---|---|---|---|
| | | | | | | | 969-974 (2015). |
| | Mouse electrophysiological study and cells | Chronic brain implant | - | Azobenzene compounds | 25-116 mW $mm^{-2}$ (473 nm, mouse) and 18-20 mW $mm^{-2}$ (470 nm, cell) | 20~200 ms | *Nat. Nanotechnol.* **15,** 296-306 (2020) |
| Magnetothermal and magnetocaloric | Anesthetized mouse and cells | Through scalp and skull | + | Nanoparticles (22 nm) | 15 kA $m^{-1}$ (500 kHz) | 5 s (cell) | *Science* **347**, 1477-1480 (2015). |
| | C. elegans behavioral study and cells | Non-invasive for C. elegans | + | Nanoparticles (6 nm) | 0.67-1 kA $m^{-1}$ (40 MHz) | 6 s (C. elegans) and 15 s (cell) | *Nat. Nanotechnol.* **5**, 602-606 (2010). |
| | Mouse behavioral study and cells | Through scalp and skull | + | Nanoparticles (15.65±2.4 nm) | 7.5 kA $m^{-1}$ (570 kHz, mouse) and 22.4 kA $m^{-1}$ (412.5 kHz, cell) | 22.8±2.6 s (mouse) and 2.18±0.17 s (cell) | *eLife* **6**, e27069 (2017). |
| | Cells | N/A | N/A | Ferritin (~12 nm) | ~275 mT (0.08 Hz) | >10 s | *Biophys. J.* **116**, 454-468 (2019). |

* '+' means 'neuron-specific'; '-' means 'nonspecific'.

**Supplementary Table 4.** Previous reports on temperature and duration threshold of thermal damage to the brain.

| **Temperature threshold** | **References** |
|---|---|
| >25 min at **42 °C** to induce thermal damage of the brain | *Int. J. Hyperther.* **19**, 267-294 (2003)<br>*Int. J. Hyperther.* **27**, 320-343 (2011) |
| No increasing sign of apoptosis or reactive microglia up to **43.3 °C** | *Science* **359**, 679-684 (2018) |
| ≥30 min at **45 °C** to induce a cytotoxic effect in neural tissue | *J. Neurochem.* **87,** 958-968 (2003) |
| Minor hyperthermic effects ($P < 0.5\%$) when exposed for 30 s at **44 °C** | *Science* **347**, 1477-1480 (2015) |

**Supplementary Table 5.** Summary of optical properties of scalp, skull and brain tissues used for Monte Carlo simulation.

| | | Optical properties | | |
|---|---|---|---|---|
| | | $\mu_a$ (mm$^{-1}$) | $\mu_s'$ (mm$^{-1}$) | $n$ |
| Scalp | 635 nm | 0.0953 | 0.945 | 1.38 |
| | 980 nm | 0.101 | 0.392 | |
| | 1064 nm | 0.0910 | 0.269 | |
| Skull | 635 nm | 0.059 | 2.68 | 1.56 |
| | 980 nm | 0.0220 | 1.80 | |
| | 1064 nm | 0.0220 | 1.60 | |
| Brain | 635 nm | 0.0267 | 0.969 | 1.35 |
| | 980 nm | 0.0614 | 0.500 | |
| | 1064 nm | 0.0233 | 0.441 | |

**Supplementary Table 6.** Summary of thermal properties of scalp, skull and brain tissues used for temperature dynamics simulation.

| | Thermal properties | | |
|---|---|---|---|
| | $C_v$ (J kg$^{-1}$ K$^{-1}$) | $k$ (W m$^{-1}$ K$^{-1}$) | $\rho$ (kg m$^{-3}$) |
| Scalp | 3391 | 0.2 | 1109 |
| Skull | 1794 | 0.41 | 1543 |
| Brain | 3630 | 0.5 | 1046 |

**Supplementary Table 7.** Antibodies used in this study.

| Primary antibodies | Secondary antibodies |
|---|---|
| Chicken anti-TH (1:1000, ab76442, abcam) | Goat anti-chicken, Alexa Fluor 647 (1:250, ab150171, abcam) |
| Rabbit anti-c-Fos (1:1000, ab222699, abcam) | Goat anti-rabbit, Alexa Fluor 488 (1:200, ab150077, abcam) |
| Mouse anti-TRPV1 (1:100, ab203103, abcam) | Goat anti-mouse, Alexa Fluor 568 (1:000, ab175473, abcam) |
| Rabbit anti-NeuN (1:200, ab177487, abcam) | Goat anti-rabbit, Alexa Fluor 488 (1:400, ab150077, abcam) |
| Rat anti-GFAP (1:500, 13-0300, Thermo Fisher Scientific) | Goat anti-rat, Alexa Fluor 647 (1:200, ab150159, abcam) |
| Mouse anti-NF (1:400, 837904, BioLegend) | Goat anti-mouse, Alexa Fluor 568 (1:200, ab175473, abcam) |
| Rabbit anti-Iba1 (1:1000, 013-27691, Wako Chemicals) | Donkey anti-rabbit, Alexa Fluor 594 (1:500, A-21207, Invitrogen) |
| Rabbit anti-Cleaved Caspase-3 (1:1000, 9664, Cell Signaling Technology) | Donkey anti-rabbit, Alexa Fluor 594 (1:500, A-21207, Invitrogen) |

**Supplementary Movie 1. Distant 1064-nm illumination induces mouse circling through intact scalp in a naturally behaving mouse.** This movie shows clockwise circling of the mouse with MINDS in the left motor cortex when illuminated with wide-field 1064-nm laser, and lack of obvious circling behavior when light is off. The frame rate is 25 frames per second (fps) and the video is played at 4× real time. Detailed protocol for MINDS injection and distant NIR-II illumination for neural modulation is described in the **Methods**.

**Supplementary Movie 2. Mouse trajectory in a Y-maze during the pretest.** This movie shows the random exploratory behavior of the mouse in the pretest, indicating no place preference before conditioning. The frame rate is 25 frames per second (fps) and the video is played at 3× real time.

**Supplementary Movie 3. Mouse trajectory in a Y-maze during the posttest.** This movie shows the strong place preference of the mouse to the left arm terminal associated with NIR-II illumination during the posttest after three consecutive days of conditioning. The frame rate is 25 frames per second (fps) and the video is played at 3× real time. Detailed protocol for MINDS injection and distant NIR-II illumination for neural modulation is described in the **Methods**.
